# Long Noncoding RNA MALAT1 is Dynamically Regulated in Leader Cells during Collective Cancer Invasion

**DOI:** 10.1101/2022.07.26.501593

**Authors:** Ninghao Zhu, Mona Ahmed, Joseph C. Liao, Pak Kin Wong

**Affiliations:** Department of Biomedical Engineering, The Pennsylvania State University, University Park, PA, USA; Department of Urology, Stanford University School of Medicine, Stanford, California, USA; Department of Mechanical Engineering and Department of Surgery, The Pennsylvania State University, University Park, PA, USA

**Keywords:** metastasis, lncRNA, bladder cancer, single cell analysis, biosensor

## Abstract

Cancer cells invade collectively with leader-follower organization. However, how leader cells are regulated during the dynamic invasion process remains poorly understood. Using a FRET nanobiosensor that tracks lncRNA dynamics in live single cells, we monitored the spatiotemporal distribution of lncRNA during collective cancer invasion. We show that lncRNA MALAT1 is dynamically regulated in the invading fronts of cancer cells and patient-derived organoids. The abundance, diffusivity, and distribution of MALAT1 transcripts are distinct between leader and follower cells. MALAT1 expression increases when a cell acquires the leader cell role and decreases when the migration process stops. Transient knockdown of MALAT1 prevents the formation of leader cells and abolishes the migration of cancer cells. Taken together, our single cell analysis suggests MALAT1 dynamically regulates leader cells during collective cancer invasion.

## Main

Collective invasion is increasingly recognized as a dominant mechanism in cancer metastasis^1^. In particular, specialized cancer cells at invading fronts of tumors, termed as ‘leader cells’, generate invasion paths, coordinate follower cells, and enhance survival in the metastatic cascade by engaging various mechanical, genetic, and metabolomic programs^2^. Genetic and epigenetic factors, stromal cells, and matrix properties have all been shown to promote the formation of leader cells and modulate their functions^3, 4, 5, 6^. Leader cells can switch positions with follower cells during invasion^7^, and ablation of leader cells initiates new leader cells^8^, underscoring the time-dependent and competitive nature of leader cell regulation. Nevertheless, how leader cells are dynamically regulated during collective cancer invasion is largely unknown.

Long noncoding RNAs (lncRNAs), which are non-protein coding RNA transcripts over 200 nucleotides, have been implicated in the progression and metastasis of cancer^9, 10^. Despite lacking the protein-coding potential, lncRNAs can control gene expression via various mechanisms, such as regulating chromatin modifying enzymes in epigenetic regulation, perturbing alternative splicing in post-transcriptional regulation, and modulating cytoplasmic RNA or protein in post-translational regulation^11, 12, 13^. Several lncRNAs, such as metastasis associated lung adenocarcinoma transcript 1 (MALAT1) and urothelial cancer associated 1 (UCA1), are reported to regulate cancer progression and metastasis^9, 14, 15, 16^. For instance, MALAT1, as the name suggests, promotes metastasis of lung and other cancer by regulating metastasis-associated genes and changing splicing patterns^17, 18, 19, 20^. In contrast, MALAT1 is also reported to suppress metastasis in colon and breast cancer by inactivating pro-metastatic transcription factors and modulating epithelial-mesenchymal transition (EMT)^21, 22, 23, 24^. These reports highlight the complex multifunctionalities of lncRNA in cancer and the challenges of studying the function of lncRNAs, calling for novel technologies that can better resolve the actions of lncRNA in cancer.

This study aims to characterize the role of MALAT1 in leader cell formation and cancer invasion. Since leader cells represent only a small subset of cancer cells, live single cell biosensors with a high spatiotemporal resolution are required for investigating the function of lncRNA^25, 26, 27, 28^. However, existing techniques, such as RNA sequencing and RNA fluorescence in situ hybridization (FISH) that lyse or fix the samples^25, 29, 30^, fail to reveal the spatial and temporal dynamics of RNAs in cancer cells during collective cancer invasion. Here, we report a fluorescence resonance energy transfer (FRET) nanobiosensor for dynamic lncRNA analysis in live single cells in 2D and 3D invasion models. Single molecule tracking was applied to determine the abundance, distribution, and dynamics of lncRNA transcripts in leader and follower cells. The expression of MALAT1 was monitored in tumor organoids derived from patients with muscle invasive bladder cancer. The formation and termination of leader cells in invading tumor structures were characterized to investigate the function of MALAT1 in collective cancer invasion.

### Visualizing lncRNA in live cancer cells with the FRET nanobiosensor

A FRET nanobiosensor was designed for characterizing lncRNA dynamics of live cancer cells at the single cell level. The nanobiosensor scheme combines the double-stranded lock nucleic acid (dsLNA) probe, the dual probe design, and FRET-based sensing to enhance the signal to noise ratio^31, 32, 33^. In particular, the nanobiosensor consists of two pairs of dsLNA probes with fluorophore and quencher sequences (**Fig. 1a**). The fluorophore labeled LNA probes (donor and acceptor) are complementary to a specific, accessible region of the target lncRNA sequence.

**Fig. 1.**
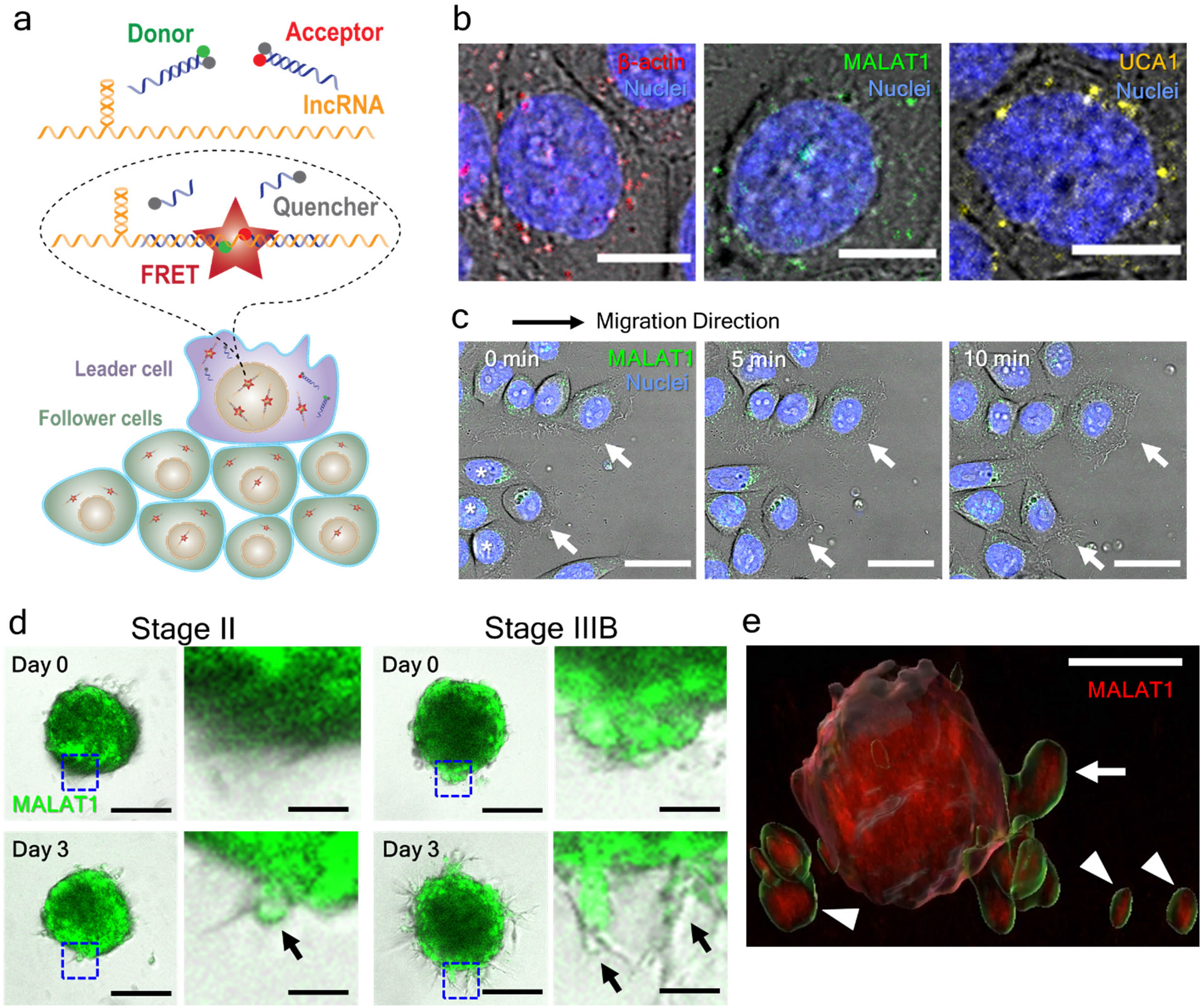
A live single cell biosensor for probing long noncoding RNA dynamics during collective cancer invasion. **a**, Schematics of the FRET-based dual double-stranded LNA nanobiosensor for studying leader and follower cells. **b**, Intracellular distributions of β-actin, MALAT1, and UCA1 RNA transcripts detected by the nanobiosensors in live cancer cells (5637). Scale bars, 10 μm. Images are representative of eight experiments. **c**, Time-lapse images for tracking MALAT1 dynamics in leader cells (marked by white arrows) during collective cell migration. Scale bars, 30 μm. Images are representative of five experiments. **d**, Detection of MALAT1 transcripts in tumor organoids derived from patients with stage II and stage IIIB muscle invasive bladder cancer. Blue dashed squares indicate the regions of the zoom-in views (right). Black arrows indicate protruding structures with leader cells sprouting from the tumor organoids. Scale bars, 200 μm (left) and 40 μm (right). Images are representative of six (Stage II) and ten (Stage IIIB) organoids. **e**, 3D rendering of leader cells (white arrows) and dissociated cancer cells (white arrowheads) invading from a cancer spheroid. Scale bar, 200 μm.

Without a target molecule, the fluorophores of donor and acceptor probes are separated and quenched, resulting in a low background signal. In the presence of a target molecule, the quencher probes are displaced, and the donor and acceptor dsLNA probes are brought to close proximity by binding to the target sequence, allowing effective energy transfer from donor to acceptor for generating a strong FRET signal (**Supplementary Information Fig. S1**). The FRET approach reduces the background noise associated with autofluorescence and thermal dissociation of the quencher probe, facilitating detection of endogenous RNA transcripts in live single cells (**Fig. 1b** and **Supplementary Information Fig. S2**). The FRET signal was validated by transient knockdown with siRNA and TGF-β1 stimulation (**Supplementary Information Fig. S3-4**). The biosensor was compatible for detecting lncRNA expressions in migrating cell monolayers, patient derived tumor organoids, and 3D cancer spheroids (**Fig. 1c-e** and **Supplementary Movie 1**).

The expressions of β-actin, MALAT1, and UCA1 transcripts were measured in live bladder cancer cells (5637). The RNA transcripts were detected by the fluorescent puncta in the FRET channel, which represent one or more RNA transcripts (**Fig. 2a-c**). The nuclear and cytoplasmic signals were examined to evaluate the distribution of RNA transcripts in live cells. A live cell nuclear dye was applied to detect the colocalization of the RNA transcripts and the nuclei by confocal microscopy. The number of transcripts of β-actin, MALAT1, and UCA1 in the cytoplasm was generally higher than the number of the transcripts colocalized with the nucleus. With TGF-β treatment, which induces EMT^34^, the total MALAT1 and UCA1 expressions were increased in the cells (**Fig. 2d**). A similar ratio of MALAT1 (but not UCA1) upregulation was observed in the nuclei (**Fig. 2e**). In contrast, the β-actin mRNA expression levels were not changed in the cytoplasmic and nuclear regions.

**Fig. 2.**
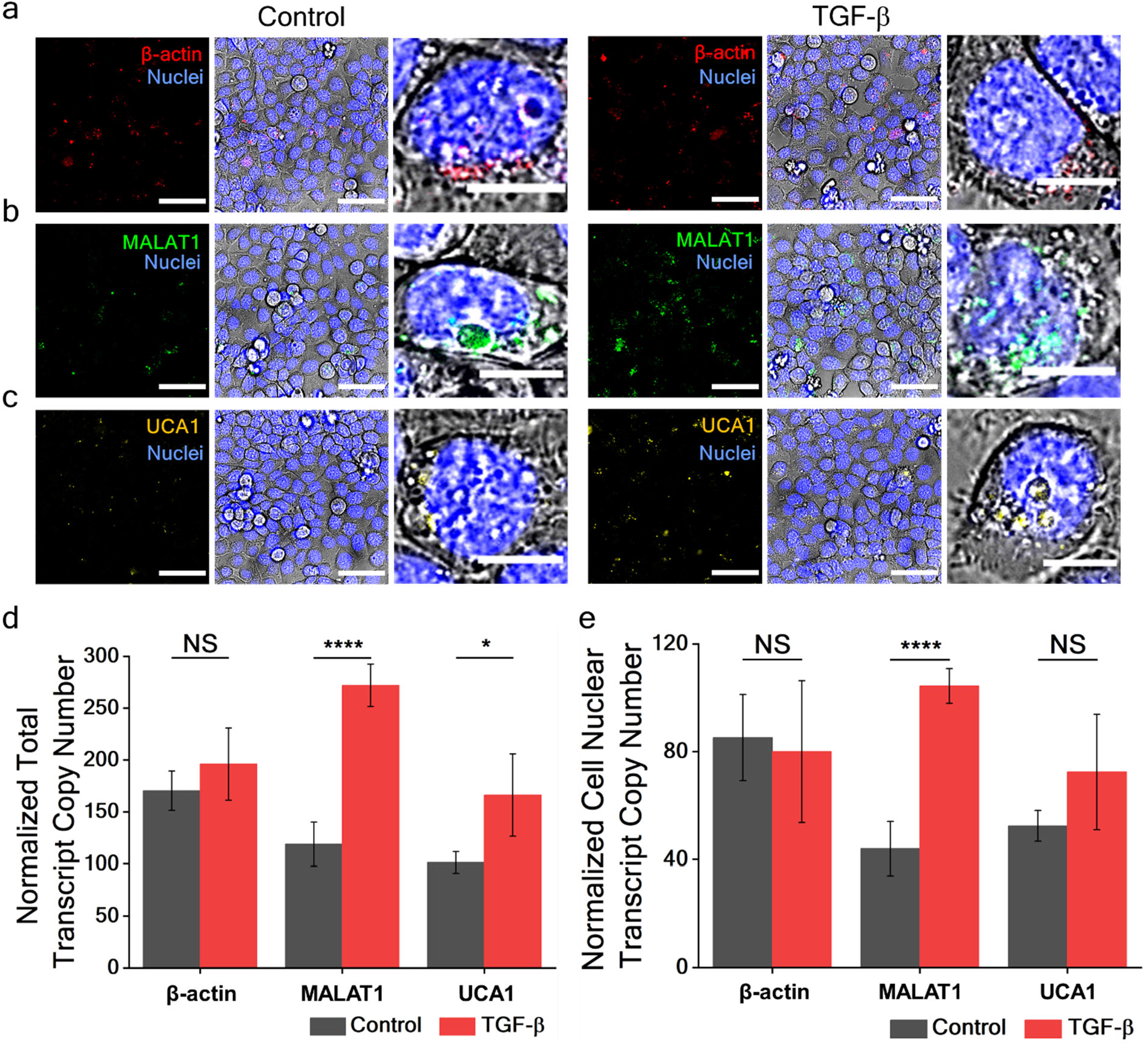
TGF-β promotes nuclear and cytoplasmic MALAT1 expression. **a-c**, Confocal images of β-actin, MALAT1, and UCA1 transcripts in live bladder cancer cells (5637). The cells were treated with buffer control and TGF-β. FRET channels (left), merged FRET and brightfield channels (middle), and zoom-in views of single cells (right) are shown to illustrate the expression distributions. Images are representative of five experiments. Scale bars, 50 μm (left and middle), 10 μm (right). **d-e**, Transcript copy numbers of β-actin, MALAT1, and UCA1 RNA (**d**) in whole cells and (**e**) colocalized with the nuclei. One-way ANOVA followed by Tukey’s post hoc test was used to compare transcript numbers between control and TGF-β treatment (n = 5, NS not significant, * p<0.05, **** p < 0.0001).

### Analyzing lncRNA distributions during collective cancer migration

The scratch cell migration assay (often called the wound healing assay) was applied to study the dynamics of lncRNA in collectively migrating cancer cells^35^. In agreement with previous studies, the cells migrated coherently and formed protruding tips with the leader-follower organization (**Fig. 3a**)^36, 37^ The formation of migration tips and leader cells was clearly observed eight hours after scratching. In this study, the cells at the protruding tips were classified as leader cells, and the cells behind were considered as follower cells. Notably, MALAT1 expression was significantly higher in leader cells compared to follower cells. On the other hand, β-actin mRNA and UCA1 expressions were comparable between leader cells and follower cells.

**Fig. 3.**
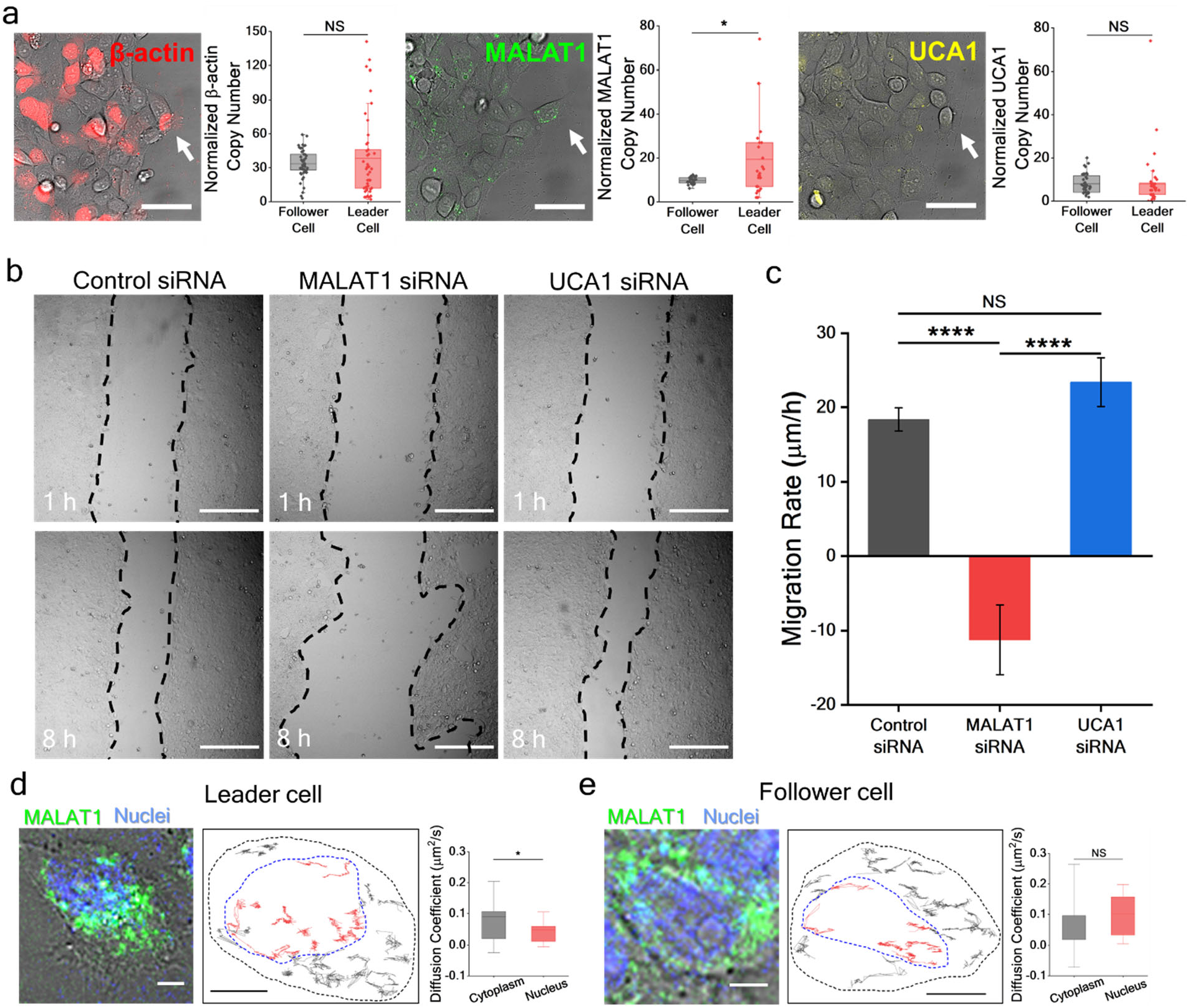
MALAT1 is upregulated in leader cells during collective cancer migration. **a**, Detection of β-actin, MALAT1, and UCA1 RNA in leader cells and follower cells. Collective cell migration was induced by scratching the cell monolayer. Cells at the protrusion tip with an aggressive morphology are considered leader cells (white arrows), and cells behind the leader cells were considered follower cells. Images are representative of four experiments. Scale bars, 50 μm. Student’s t-test was used to compare the transcript numbers between the leader and follower cells (n ≥ 25, NS not significant, * p < 0.05). **b**, Collective migration of cancer cells treated with control, MALAT1 siRNA, and UCA1 siRNA at 1 hour and 8 hours. Black dotted lines illustrate the boundary of the migrating cells. Images are representative of four experiments. Scale bars, 300 μm. **c**, Cell migration rate estimated by the average distance between the cell boundaries. Oneway ANOVA followed by Tukey’s post hoc test was used to compare the migration rates (n = 4, NS not significant, **** p < 0.0001). **d-e**, Single molecule tracking of MALAT1 transcripts in leader cells and follower cells. (Left) Representative images from eight experiments. Scale bars, 5 μm. (Middle) Traces of MALAT1 transcripts. Black and blue dashed lines indicate the cell boundary and the nucleus, respectively. Scale bars, 5 μm. (Right) Comparison of cytoplasmic and nuclear diffusivities of MALAT1. Student’s t-test was used to compare the diffusivities of MALAT1 RNAs in the nucleus and cytoplasm (n ≥ 5, NS not significant, * p < 0.05).

The functions of MALAT1 in collective cancer migration were investigated by siRNA treatment (**Fig. 3b-c**). Transient knockdown of MALAT1 abolished the formation of migration tips. The boundary of the monolayer retracted slightly and resulted in a negative average migration rate. Examining the cell morphology revealed a non-migratory phenotype. In contrast, downregulation of UCA1 did not display a substantial change in the cell morphology, and the migration rate was comparable to the control siRNA. We also treated the migrating monolayers with TGF-β (**Supplementary Information Fig. S5**). While coherent monolayers were observed in the control group, TGF-β treatment disrupted the cell-cell adhesion and dissociated cancer cells in the scratched monolayers. MALAT1 expression was enhanced in some dissociated cells.

We further examined MALAT1 in leader and follower cells by single molecule tracking (**Fig. 3d-e** and **Supplementary Movies 2-3**). The mean squared displacement and diffusivity of the molecules were estimated by tracking the position of transcripts as a function of time. The diffusivity of cytoplasmic MALAT1 was approximately 0.09 μm^2^/s, which is comparable with β-actin mRNA.^32, 38^ Interestingly, we observed a reduced MALAT1 diffusivity in the nuclei of leader cells. In leader cells, MALAT1 in the cell nuclei exhibited a lower diffusivity (0.050 ± 0.011 μm^2^/s) compared to MALAT1 in the cytoplasm (0.089 ± 0.017 μm^2^/s). In contrast, the MALAT1 diffusivity in follower cells was similar between the nuclei (0.010 ± 0.043 μm^2^/s) and cytoplasm (0.097 ± 0.033 μm^2^/s). We also observed a portion (~50%) of MALAT1 transcripts was immobile (< 0.005 μm^2^/s); however, the portion was similar between leader and follower cells (**Supplementary Fig. S6**). The lower diffusivity suggests MALAT1 in leader cells may engage in different molecular complexes and activities compared to follower cells. These observations reveal that MALAT1 is differentially activated in leader cells during collective cancer migration.

### Dynamics of MALAT1 in 3D collective invasion

We further evaluated the relationship between MALAT1 and leader cells using a 3D tumor invasion assay^37, 39^. We formed 3D cancer spheroids with bladder cancer cells transfected with the FRET nanobiosensors. The nanobiosensor was transfected into the cells before spheroid formation to ensure uniformity of probe transfection. The tumor spheroids were embedded in an invasion matrix consisting of collagen I and basement membrane extract, representing the lamina propria and basement membrane compositions. TGF-β (40 ng/ml) was added to the invasion matrix to induce invasion. The cancer spheroids were allowed to invade for a period of three days (**Fig. 4a**). Protruding cells were observed in approximately 24 hours. The sprouting structures continued to grow and invaded into the invasion matrix. Examining the spatial distribution revealed that MALAT expression was upregulated at the outer region of the spheroid, where the cancer cells interfaced with the invasion matrix. Interestingly, as the sprout continued to invade the matrix, the MALAT1 expression of the sprouts also increased during the three-day invasion period. As a control, β-actin mRNA decreased slightly in the same period (**Fig. 4b**).

**Fig. 4.**
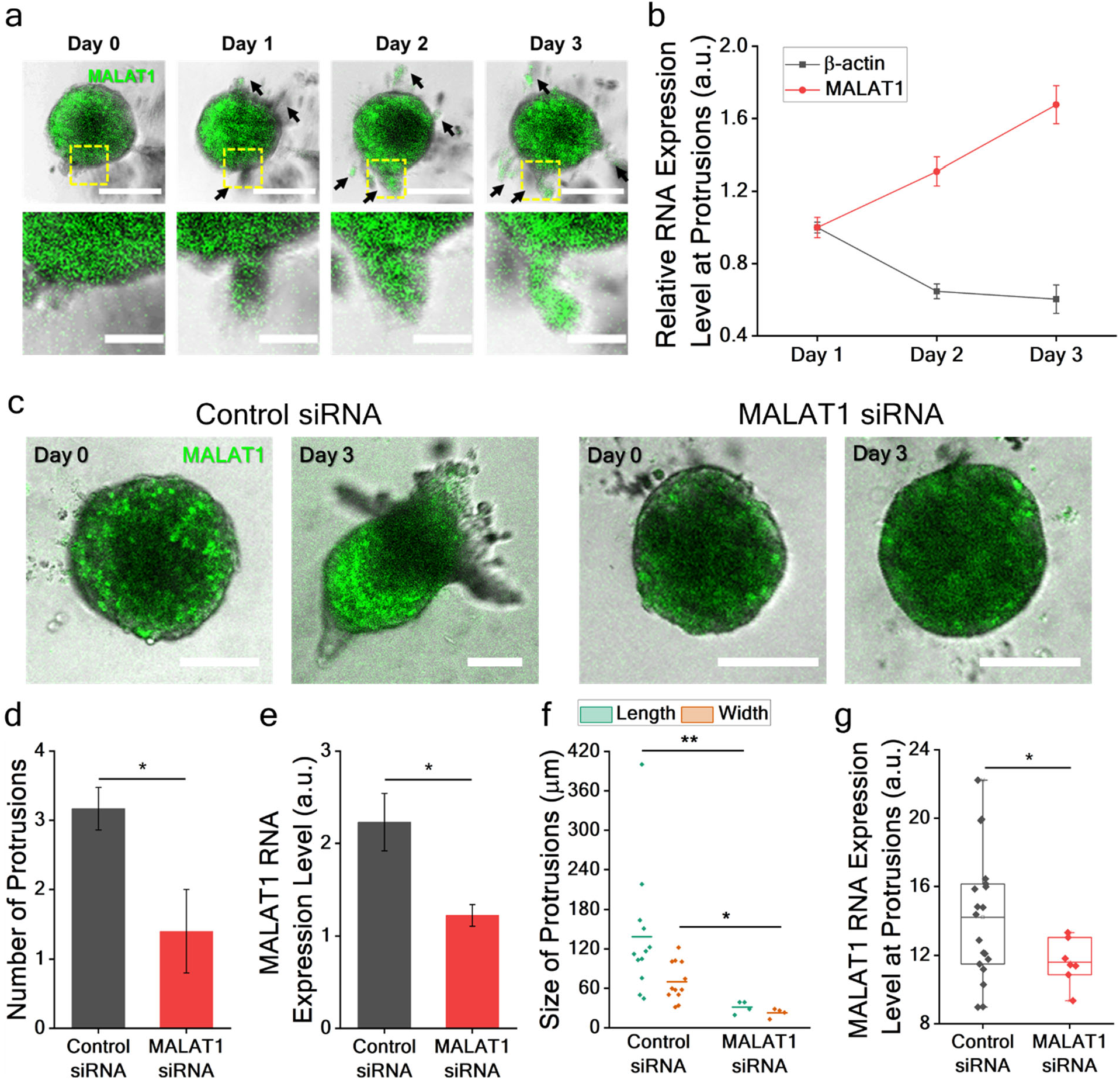
MALAT1 is upregulated in invading sprouts of 3D bladder cancer spheroids. **a**, Bladder cancer spheroids (5637) were treated with TGF-β and allowed to invade the invasion matrix for three days. Black arrows indicate invading protrusions from the spheroid. Yellow dashed squares highlight a protruding structure in the zoom-in views (bottom). Scale bars, 250 μm (top) and 80 μm (bottom). Images are representative of five experiments. **b**, The expression of β-actin mRNA and MALAT1 at protrusions. The value was normalized to the initial intensity. One-way ANOVA was used to analyze the RNA expression level, indicating an increasing MALAT1 expression (p = 5.82×10^-5^) and a decreasing β-actin expression (p = 0.0023) (n ≥ 3). **c**, MALAT1 and control siRNA treated spheroids in the 3D invasion assay. Scale bars, 150 μm. Images are representative of three experiments. **d**, Comparison of the number of protrusions per spheroid with control siRNA and MALAT1 siRNA on day 3. **e**, Comparison of MALAT1 expression of cancer spheroids with control and MALAT1 siRNA. Student’s t-test was used to compare the number of protrusions and the MALAT1 expression (n = 6 for control siRNA and n = 5 for MALAT1 siRNA, *p < 0.05). **f**, The length and width of protrusions from the cancer spheroids on day 3. Two-way ANOVA followed by Tukey’s post hoc test was used to compare the dimensions of protrusions from spheroids with MALAT1 siRNA and control siRNA (n = 4 for MALAT1 siRNA and n = 12 for control siRNA, * p < 0.05, **, p < 0.01). **g**, MALAT1 expression level at the protrusions of spheroids treated with MALAT1 siRNA and control siRNA on day 3. Student’s t-test was used to compare the expression. (n = 7 for MALAT1 siRNA and n = 19 for control siRNA, * p < 0.05).

We knocked down MALAT1 by siRNA to investigate the role of MALAT1 in 3D collective cancer invasion (**Fig. 4c**). Similar to the 2D experiment, the formation of invasive sprouts and leader cells was attenuated by MALAT1 siRNA (**Fig. 4d**). The FRET biosensor revealed that the cancer spheroids with MALAT1 siRNA had a lower MALAT1 expression level (**Fig. 4e**). The length and width of the protrusions from the cancer spheroids were decreased in the MALAT1 siRNA group (**Fig. 4f** and **Supplementary Information Fig. S7**). The results suggest the number of follower cells invading the matrix were attenuated. In both control and MALAT1 siRNA, MALAT1 was upregulated near the outer surface of the spheroids. At the protrusions, the MALAT1 expression was reduced in the MALAT1 siRNA group compared to the control group (**Fig. 4g**). These results indicate MALAT1 is associated with the formation of leader cells and collective invasion in the 3D microenvironment.

### MALAT1 in human bladder cancer samples

We evaluated the involvement of MALAT1 in patient derived cancer cells and tumor organoids^40^. The cancer cells were dissociated from tumor samples from patients diagnosed with stage II (spread into the bladder muscle wall), stage IIIB (spread to two or more local lymph nodes), and stage IV (spread to distant lymph nodes and organs) urothelial carcinoma. Single cell analysis with the FRET nanobiosensor revealed that MALAT1 was highly heterogeneous among the cancer cells. Specifically, a subset (5-10%) of cells expressed a high level of MALAT1 that was visually distinct from the rest of the population (**Fig. 5a**). The heterogeneity could not be explained by the transfection efficacy between the primary cancer cells, as transfection was effective and relatively uniform for detecting β-actin mRNA. We analyzed the MALAT1 expression and estimated the portion of MALAT1 expressing cells. The MALAT1 expression in the stage IIIB sample was significantly higher than the value in the stage II sample. In contrast, the β-actin mRNA expressions were similar in the samples (**Fig. 5a**). Nevertheless, the portions of MALAT1 cells were similar between the stage II and stage IIIB samples, suggesting the increase was mainly contributed by an enhanced MALAT1 expression instead of the portion of MALAT1 expressing cells.

**Fig. 5.**
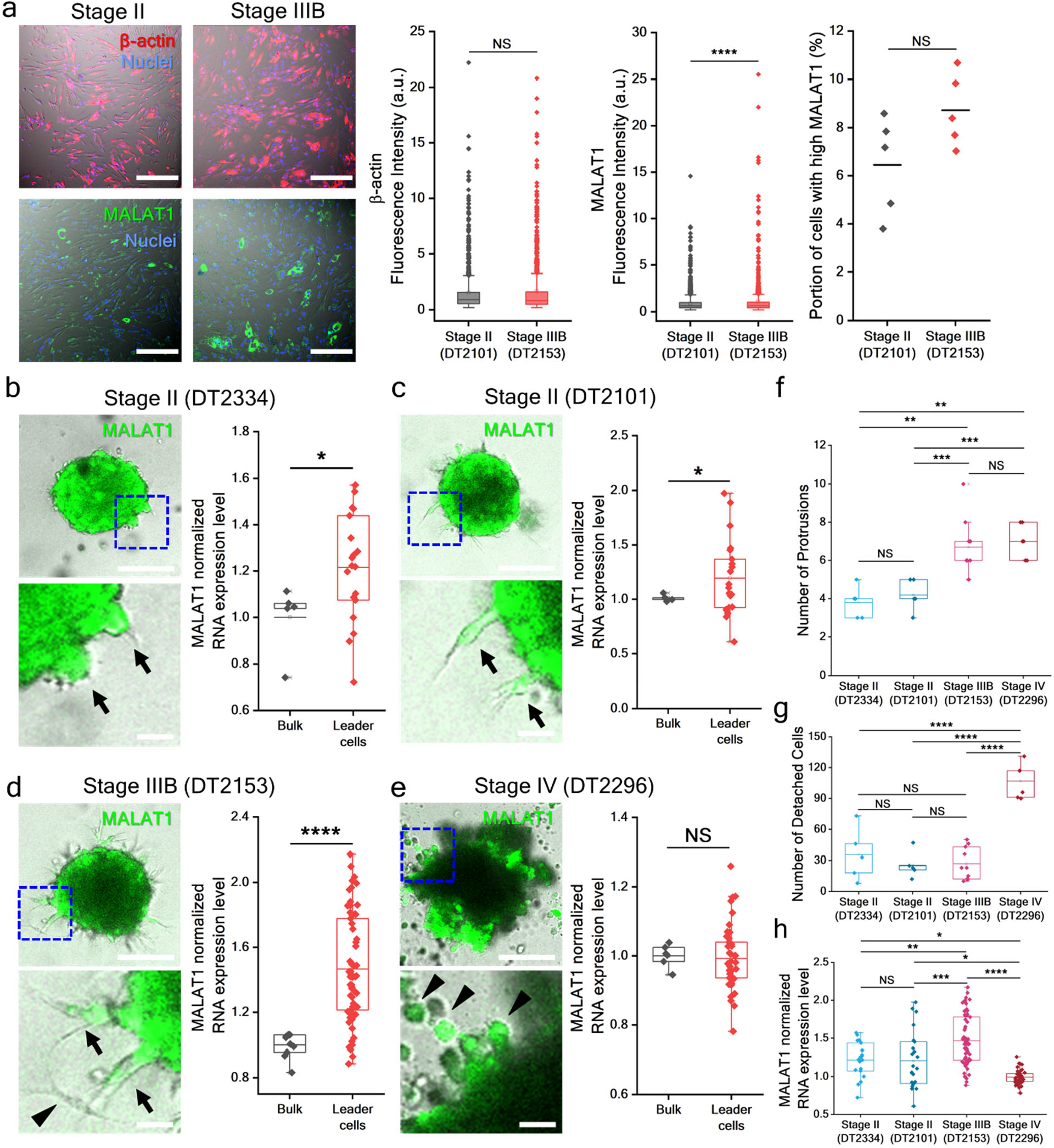
MALAT1 is associated with the stage and invasiveness of bladder cancer. **a**, β-actin mRNA and MALAT1 expressions in cancer cells derived from bladder cancer patients. Scale bars, 300 μm. MALAT1 is higher in the stage IIIB sample compared to the stage II sample. Student’s t-test was used to compare between samples (n ≥ 1148 for gene expression and n = 5 for cell portions with an enhanced MALAT1 expression). **b-e**, MALAT1 expression in 3D human tumor organoids derived from bladder cancer patients on day 3. Blue dashed squares highlight the zoom-in views (bottom) with leader cells (black arrows) or dissociated cells (black arrowheads). Scale bars, 200 μm (top) and 100 μm (bottom). Images are representative of 5 (DT2334 and DT2101), 6 (DT2296), and 10 (DT2153) organoids. Student’s t-test was used to compare MALAT1 expression (n = 5, 5, 10, and 6 for organoids of DT2334, DT2101, DT2153, and DT2296, n = 19, 23, 67, and 42 for leader cells of DT2334, DT2101, DT2153, and DT2296). **f**, The number of protrusions per organoid on day 3. One-way ANOVA followed by Tukey’s post hoc test was used to compare the number of protrusions from different samples (n = 5, 5, 10, and 6 for DT2334, DT2101, DT2153, and DT2296). **g**, The number of detached cells per organoid on day 3. One-way ANOVA followed by Tukey’s post hoc test was used to compare the number of protrusions from different human samples (n = 5, 5, 10, and 6 for DT2334, DT2101, DT2153 and DT2296). **h**, Normalized MALAT1 expression in leader cells. One-way ANOVA followed by Tukey’s post hoc test was used to compare the MALAT1 expression in leader cells between samples (n = 19, 23, 67, and 42 for DT2334, DT2101, DT2153 and DT2296, NS, not significant, *, p < 0.05, **, p < 0.01, ***, p < 0.001, ****, p < 0.0001).

The patient derived cancer cells were also studied using the scratch cell migration assay (**Supplementary Information Fig. S8**). However, the cell monolayer was less coherent compared to typical epithelia in wound healing studies^37, 41^. After scratching, the cells exhibited an elongated, spindle-shaped morphology near the migrating front. While leader-follower organization could still be observed in some cases, the cells were not physically aligned, suggesting a weak physical interaction. Furthermore, we did not observe a substantial change in the number of MALAT1 expressing cells. MALAT1 expression was randomly expressed in leader and follower cells in the loose monolayer.

We investigated the function of MALAT1 by generating tumor organoids with the patient derived cancer cells and performed a 3D invasion assay (**Fig. 5b-e** and **Supplementary Information Fig. S9)**. For all patient derived samples tested, MALAT1 was strongly expressed at the surface of the tumor organoids. Since the probes were transfected into the cells before forming 3D organoids, the spatial distribution was unlikely a result of probe accessibility in 3D culture. Examining the β-actin control also supported the spatial distribution could not be fully explained by optical penetration. Sprouting invasion and cell detachment were observed from the organoids (**Fig. 5f-g**). Protruding structures with the leader-follower organization could be observed at the surface of the organoids. The number of protrusions increased with the clinical stage. For stage II and IIIB samples, coherent invading sprouts with clear leader-follower organization and filopodia. In contrast, protrusions from the stage IV organoids showed a loose, irregular morphology. The cells also exhibited a round shape without any apparent filopodial structures. There were many detached cells in the stage IV organoids compared to other organoids. The stage IV sample appeared to engage in a distinct invasion mechanism compared to other samples. Moreover, examining the invading sprouts and detached cells revealed that MALAT1 was enhanced in the stage IIIB sample compared to the stage II samples (**Fig. 5h** and **Supplementary Information Fig. S10**). These results further support MALAT1 is involved in coordinating the collective invasion of bladder cancer.

### Real-time dynamics of lncRNA in leader cells

Since our results suggest MALAT1 expression is associated with cell coordination, we performed single cell tracking experiments during collective cancer migration. Time-lapse images of the migrating monolayer were taken near the leading edges during cell migration and closure of the monolayer (**Fig. 6a** and **Supplementary Movie 4**). The expressions of MALAT1 were dynamically monitored in cells near the migrating front. The nuclear MALAT1 transcript was analyzed to study the MALAT1 intracellular distribution and compare with the cell behaviors. **Fig. 6b** shows leader cell switching and termination during collective cell migration (see also **Supplementary Movies 5** and **Supplementary Information Fig. S11**). A follower cell (marked by yellow dash lines) migrated to the leading edge (0-30 minutes) and took over the leader cell position. The cell also acquired an aggressive morphology with active lamellipodia (30-60 minutes). The newly formed leader cell led follower cells and merged with the other leading edge. The leader cells contacted the other leading edge at 70 minutes (i.e., the cell-contact time), and the edges continued to merge (120 minutes). Remarkably, the portion of nuclear MALAT1 strongly correlated with the position of the cells. In particular, the nuclear MALAT1 remained steady at the basal level (~30%) from 0-30 minutes when the cell was in the follower position. From 30-60 minutes, when the cell took over the leader position and exhibited an aggressive morphology, the nuclear MALAT1 increased from 30% to over 60%. From 60 min to 120 min, when the leading edges merged to form a monolayer, the nuclear MALAT1 returned to the basal level gradually (most rapidly between 60-90 minutes).

**Fig. 6.**
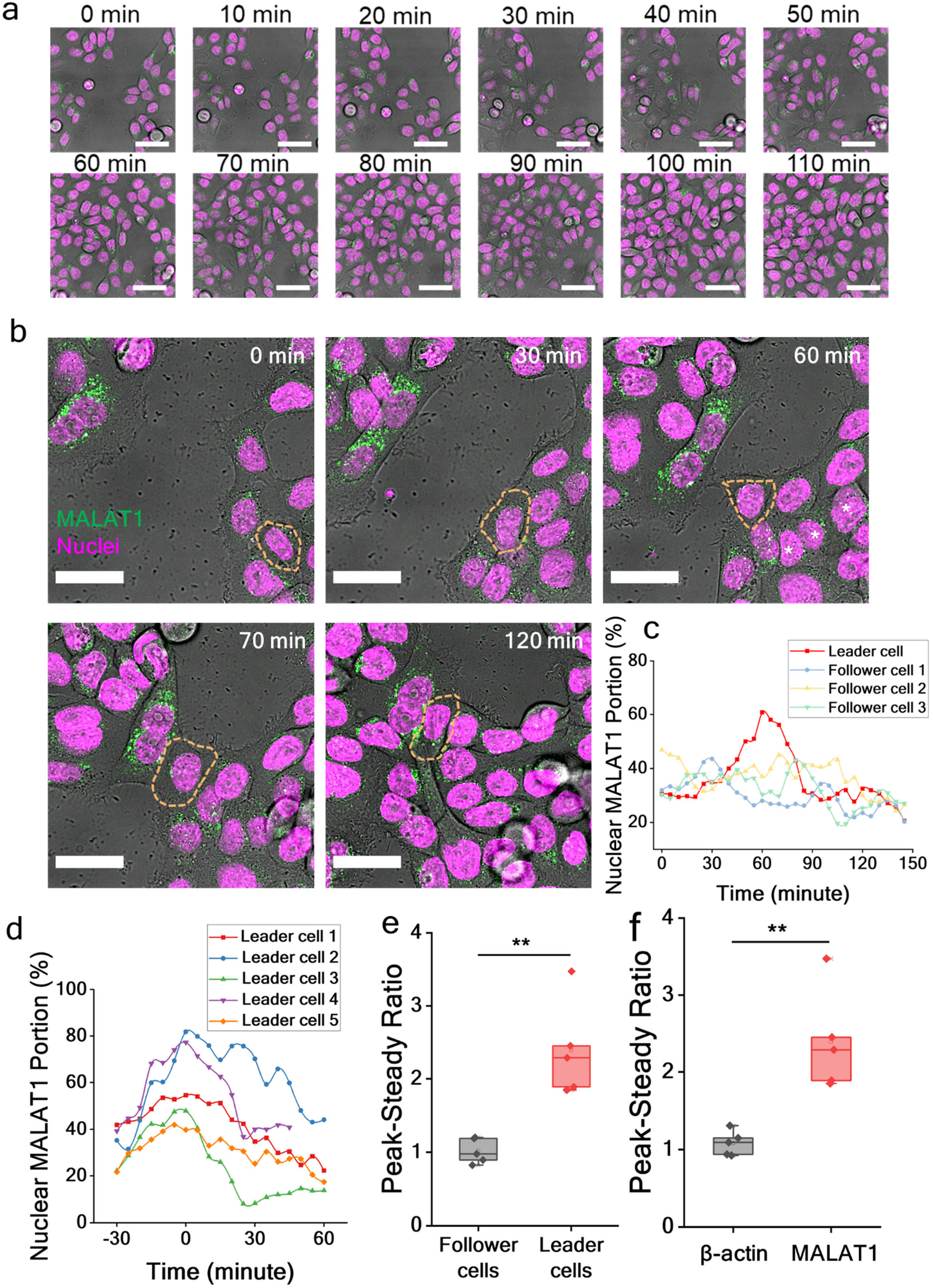
Dynamics tracking of MALAT1 during the formation and termination of leader cells. **a**, Time lapse images of cancer cells (5673) during collective cell migration induced by the scratch assay. Scale bars, 50 μm. **b**, Time-lapse images tracking the switching and end of a leader cell (yellow dashes) near the migrating front. A follower cell was initially behind the cell boundary. The cell migrated to the front, acquired the leader cell role, and displayed an aggressive morphology. Then, the leader cell reached the other boundary and was surrounded by other cells in the monolayer. Scale bars, 20 μm. **c**, Tracking of nuclear MALAT1 of the leader cell and follower cells outlined in (**b**). Data represent the portion of nuclear MALAT1 in the cell. MALAT1 increased when the cell assumed the role of leader cell and decreased when the boundary merged. **d**, Examples of nuclear MALAT1 in leader cells during the merging of cell boundaries. Time zero (or peak) is when the leader cells reached and contacted the other boundary. MALAT1 return to the basal level (or steady state) in 30-60 minutes. **e**, Peak-steady ratio of nuclear MALAT1 in leader cells and follower cells. **f**, Peak-steady ratio of nuclear MALAT1 and β-actin mRNA in leader cells. Student’s t-test was used to compare the Peak-steady ratio (n = 5, ** p < 0.01).

We further analyzed leader cell behaviors during other cases of cell monolayer closure. When the cell-contact time was considered as time zero, a similar pattern of the nuclear MALAT1 was generally observed in leader cells (**Fig. 6d**). The nuclear MALAT1 in leader cells dropped approximately 50-60% (or a peak-steady ratio of >2) from the peak value (**Fig. 6e**). In contrast, the nuclear MALAT1 maintained constant (i.e., a peak-steady ratio of ~1) in follower cells (**Fig. 6e**). Furthermore, the change in the nuclear MALAT1 was not observed in β-actin mRNA of leader cells (**Fig. 6f**). These observations collectively supported the notion that MALAT1 is associated with leader cells and dynamically coordinates collective cancer invasion.

## Discussion

In this study, we report a FRET nanobiosensor for probing lncRNA dynamics in live single cells. We demonstrate spatiotemporal analysis of lncRNA during collective cancer invasion, which is challenging for conventional RNA detection techniques. The lncRNA biosensor is compatible with both 2D collective cell migration and 3D cancer invasion. Compared to other live cell RNA sensing techniques^25, 26^, the nanobiosensor detects endogenous RNA transcripts and is capable of detecting lncRNA in patient derived cancer cells and tumor organoids. The combination of LNA modification, the double-stranded probe, and the FRET dual probe design provides an outstanding stability, sensitivity, and signal-to-noise ratio for tracking lncRNA transcripts in live cells. The ability to monitor the abundance, diffusivity, and distribution can be useful for studying the diverse modes of action of lncRNA^11, 12, 13^.

Our results reveal that MALAT1 is associated with collective invasion of bladder cancer. MALAT1 expression was upregulated in leader cells and correlated with the aggressiveness of cancer cells. Attenuating MALAT1 disrupted collective cancer migration and reduced the formation of invading sprouts in the 3D invasion assay. As demonstrated in cancer cells derived from patients with muscle invasive bladder cancer, the invasiveness of tumor organoids in the 3D invasion matrix increased with the tumor stage. The invasiveness is indicated by the number of sprouts and the number of detached cells. The expression of MALAT1 correlates with the formation of leader cells and protrusions (but not the less coordinated stage IV sample). Notably, a single cell analysis technique is required to resolve the MALAT1 expressing leader cell population as a standard bulk RNA measurement technique (e.g., qRT-PCR) may dilute the increase in lncRNA expression in leader cells. Thus, the combination of the organoid invasion assay and the nanobiosensor targeting MALAT1 may provide a novel prognostic approach for identifying muscle invasive bladder cancer. Moreover, the stage IV (metastatic) sample displayed less coherent organoids with a large number of detached cells. The cells appeared to engage in solitary invasion and showed a spherical shape in the 3D environment, suggesting the cells are not adopting a collective invasion mechanism^1^. Future clinical studies using patient derived tumor samples (e.g., from transurethral resection of bladder tumor) are warranted for validating the translational potential of MALAT1 in bladder cancer prognosis.

Our study revealed that MALAT1 is dynamically regulated in leader cells. The mechanistic interactions between MALAT1 and leader cells have not been established. Previous studies suggest that MALAT1 modulates gene expression and interacts with various transcription factors (e.g., TEAD), microRNAs (e.g., miR200 and miR-125b), splicing factors (e.g., SR proteins), and epigenetic regulators (e.g., EZH2)^16^. The single molecule tracking feature of the nanobiosensor revealed a decrease in the nuclear MALAT1 in leader cells. The distribution of MALAT1 may contribute to the modulation of several signaling pathways associated with collective cancer invasion. In particular, MALAT1 has been reported to modulate EMT by sponging miRNA (miR200 and miR125b) and inhibiting Ezh2-Notch1 signaling^20, 42, 43, 44^. In agreement, we observed upregulation of MALAT1 with TGF-β in 5637 cells. The cells exhibited aggressive morphology, and MALAT1 transcripts were enhanced in the nuclei after TGF-β treatment. MALAT1 expression was also increased near the migrating front in the scratch cell migration assay^35^ and TGF-β induced 3D invasion^34^. Our recent study revealed that collective invasion and leader cell formation can be spatially coordinated by the Nrf2-EMT-Notch1 circuit^37^. The function of MALAT1 in modulating EMT may provide a mechanism for regulating the leader cell formation. Furthermore, MALAT1 is known to regulate cell-cell and cell-matrix interactions through the Hippo-YAP/TEAD signaling^44, 45^. As observed in the single cell tracking experiments, MALAT1 increased when a cell took over the leader position and when an invading sprout penetrated the 3D invasion matrix. In contrast, MALAT1 was downregulated when leader cells were in contact with other cells during the closure of the monolayer, and disruption of MALAT1 abolished the collective cancer invasion. These observations support that MALAT1 is dynamically regulated during interactions with the environment and neighboring cells, which are major functions of leader cells^2^. Future studies with other molecular characterization techniques, specific gene editing approaches, and physiologically relevant models will be required to elucidate the molecular mechanisms and functions of MALAT1 in leader cell formation and cancer invasion.

## Conclusion

This study demonstrates a FRET nanobiosensor with a high spatiotemporal resolution for investigating lncRNA dynamics in live single cells. Using the nanobiosensor, our results suggest a functional role of MALAT1 in the dynamic regulation of leader cells during collective cancer invasion. The finding may create new opportunities in developing novel prognostic and therapeutic approaches for cancer in the future.

## Materials and Methods

### Cell culture and reagents

The human bladder cancer cell line 5637 from American Type Culture Collection (Manassas, VA) was cultured in RPMI 1640 medium supplemented with 10% fetal bovine serum and 10 μg/mL Gentamicin (Fisher Scientific, Hampton, NH). The human bladder cancer dissociated tumor cells (stage II, lot number: DT02101 and DT02334; stage IIIB, lot number: DT02153; stage IV, lot number: DT02296) purchased from BioIVT (Westbury, NY) were cultured in Dulbecco’s Modified Eagle’s Medium (DMEM; Fisher Scientific, Hampton, NH) supplemented with 10% fetal bovine serum and 10 μg/mL Gentamicin. The cells were cultured at 37 °C in a humidified incubator with 5% CO2 and seeded in 24 well glass bottom plates (Cellvis; Mountain View, CA) 24 hours prior to experiments.

The MALAT1 siRNA (Silencer® Select, Catalog #: 4392420), UCA1 siRNA (Silencer® Select, Catalog #: 4392420) and control siRNA (Silencer® Select Negative Control No. 1 siRNA, Catalog #: 4390843) were purchased from Thermo Fisher Scientific (Waltham, MA). To knockdown MALAT1 and UCA1 expressions, 20 nM siRNAs were transfected into the cells by Lipofectamine RNAiMAX according to the manufacturer’s protocol (Fisher Scientific, Hampton, NH) for 48 hours. To modulate MALAT1 and UCA1 expressions, transforming growth factor beta 1 (TGF-β1; Bio-Techne, Minneapolis, MN) was added to the cells with a 10 ng/mL concentration for 48 hours. Cell nuclei were stained by NucSpot® Live 650 Nuclear Stains (Biotium, Fremont, CA) for 10 minutes according to the manufacturer’s protocol. Other reagents were purchased from Fisher Scientific (Hampton, NH) unless specified otherwise.

### FRET nanobiosensor design and probe transfection

Dual double-stranded locked nucleic acid (dsLNA) probes and synthetic targets were synthesized by Integrated DNA Technologies (San Diego, CA). The FRET nanobiosensor consists of two dsLNA probes, modified with a donor and an acceptor (ATTO488-Cy3 or FAM-Texas Red). The fluorophores were labeled at 5’ end of the donor and 3’ end of the acceptor (**Supplementary Tables 1-3**). The secondary structures of the lncRNAs were predicted by the mfold web server (http://www.unafold.org/)^46^ to determine the accessible region (i.e., loops) of the lncRNA. The dual probes target two adjacent sequences (separated by four nucleotides) in an accessible region of the target lncRNA. The probes were designed to be in close proximity while preventing steric hindrance. A quencher probe was designed for each fluorescence probe to silence the unbound probes and enhance the signal-to-noise ratio. All sequences were aligned to the human transcriptome by the NCBI Basic Local Alignment Search Tool (BLAST) to verify and avoid binding to non-specific RNAs.

The fluorophore and quencher probes were dissolved in sodium chloride-tris-EDTA (10 mM tris, 1 mM EDTA, and 100 mM NaCl) and were mixed at a 1:3 ratio. The probes were then incubated at 37 °C for 15 minutes to form dsLNA. The FRET-based biosensor consists of a donor dsLNA and an acceptor dsLNA mixed at a 1:4 ratio. Then, the FRET-based biosensor was transfected into the cells cultured in 24 well glass bottom plates by Lipofectamine 3000 according to the manufacturer’s protocol (Fisher Scientific, Hampton, NH). The final concentration of the donor probe in each well was 20 nM. The cells were incubated with the biosensor for 24 hours, followed by rinsing for three times with 1× phosphate-buffered saline (PBS).

### Cell migration assay

A scratch cell migration assay was performed to study collective cell migration. After reaching 100% confluency, the cell monolayer was scratched using a sterilized 200 μL pipet tip to create a cell-free region. The cell monolayer was rinsed with 1× PBS before and after scratching. The scratched cell monolayers were allowed to migrate and were imaged after eight hours. The distance between the two leading edges (wound width) was measured by Fiji ImageJ, and the migration rate was calculated by the difference of the initial and final wound width divided by migration time.

### 3D invasion assay and patient derived organoids

A 3D invasion assay was applied to investigate the lncRNA dynamics in collective cancer invasion with Cultrex© 3D Spheroid Cell Invasion Assay Kits according to manufacturer’s instructions (R&D Systems, Minneapolis, MN). Briefly, bladder cancer cell line 5637 or the human bladder cancer dissociated tumor cells with the biosensor were incubated in Spheroid Formation Extracellular Matrix in a 96 well round bottom plate (10,000 cells per cell) for three days to form spheroids. The Invasion Matrix, consisting of collagen I and basement membrane extract, was added to the spheroids and organoids. After gel formation for an hour, fresh culture media were added. The spheroids and organoids were then imaged at 0, 24, 48 and 72 hours.

### Image acquisition and data analysis

Images were obtained with a laser scanning confocal microscope (Leica TCS SP8; Leica Microsystems, Wetzlar, Germany) with an enclosed incubator (OKOlab, Italy) for temperature, humidity, and gas control. The donor probes were excited with a 488 nm solid-state laser, the acceptor probes were excited with a 514 nm (Cy3) or 552 nm (Texas Red) solid-state laser, and the cell nuclei stain was excited with a 638 nm solid-state laser. The fluorescence signals were collected with an NA = 0.9, 63× air objective. The fluorescence signals acquired from the wavelength 500 nm to 530 nm were determined as donor channel (excited with 488 nm laser) and the fluorescence signals acquired from the wavelength 530 nm (Cy3) or 570 nm (Texas Red) to 630 nm were determined as FRET channel (excited with 488 nm laser) and acceptor channel (excited with 514 nm or 552 nm laser).

For 2D monolayer experiments, a MATLAB (MathWorks, R2021a) program was developed to segment and quantify the puncta from the FRET channel of fluorescence images. The number of the puncta was used to estimate the total copy number of RNA and the portion of nuclear RNA. Images were processed by Fiji ImageJ.

For single molecule tracking, the fluorescence signal from the biosensor, representing the RNA molecule, was captured at a frame rate of 1 Hz for 5 minutes. The tracking of transcripts was processed with Fiji ImageJ plugin “TrackMate”. The DoG (differences of gaussian) detector was applied to identify the single particles with an estimated object diameter of 1.5 μm and quality threshold of 0.2. The Linear Assignment Problem (LAP) tracker was applied to detect the trajectory of each single particle. To link the particles from frame to frame, the maximum distance was set to be 1 μm. The track segments were connected when the maximum distance of 2 μm and the maximum of two gaps were allowed for each track. The artifactual tracks were filtered by discarding the tracks with less than 60 particles per track and track mean speed higher than 1.37 μm/s. To interpret the single transcript tracking, the mean squared displacement (MSD) of each track was calculated by 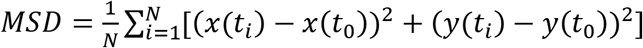, where N is the total number of particles in the track, and *x*(*t_i_*) and *y*(*t_i_*) represent the position of the particle at frame i. The MSD was then modeled by anomalous diffusion, described by *MSD* = 4*Dt^α^*, where *D* is the diffusion coefficient, *t* is the time, and *α* is the exponent^47, 48^. The diffusion coefficient of each track was acquired by the MATLAB curve fitting function.

The 3D spheroid images were analyzed by Imaris (Bitplane Inc., version 9.8). The 3D spheroids were rendered and segmented from the FRET channel of the z-stack images, and the lncRNA expression was determined by the mean fluorescence intensity.

## Statistical analysis

Data obtained from Fiji ImageJ, MATLAB and Imaris were analyzed with Origin software (OriginLab, version 9.9) and presented as mean ± standard error of mean. All experiments were performed at least three times. Student’s t-tests were used to compared two experimental groups. One-way or two-way analysis of variance followed by Tukey’s post hoc test was used to compare multiple experimental groups. The statistical significance was symbolized by NS (p > 0.05), * (p ≤ 0.05), ** (p ≤0.01), *** (p ≤0.001), or **** (p ≤0.0001).

## Acknowledgements

The authors thanks Ian Eder and Sam Vilchez for technical support of the 3D invasion assay. This work was supported by National Science Foundation (1802947 and 2033673) and NIH (R21AR079095).

## References

1. Friedl P, Locker J, Sahai E, Segall JE. Classifying collective cancer cell invasion. Nat Cell Biol 2012, 14(8): 777–783.

2. Vilchez Mercedes SA, Bocci F, Levine H, Onuchic JN, Jolly MK, Wong PK. Decoding leader cells in collective cancer invasion. Nat Rev Cancer 2021, 21(9): 592–604.

3. Wolf K, Friedl P. Extracellular matrix determinants of proteolytic and non-proteolytic cell migration. Trends Cell Biol 2011, 21(12): 736–744.

4. Zoeller EL, Pedro B, Konen J, Dwivedi B, Rupji M, Sundararaman N, et al. Genetic heterogeneity within collective invasion packs drives leader and follower cell phenotypes. J Cell Sci 2019, 132(19): jcs231514.

5. Westcott JM, Prechtl AM, Maine EA, Dang TT, Esparza MA, Sun H, et al. An epigenetically distinct breast cancer cell subpopulation promotes collective invasion. J Clin Invest 2015, 125(5): 1927–1943.

6. Gaggioli C, Hooper S, Hidalgo-Carcedo C, Grosse R, Marshall JF, Harrington K, et al. Fibroblast-led collective invasion of carcinoma cells with differing roles for RhoGTPases in leading and following cells. Nat Cell Biol 2007, 9(12): 1392–1400.

7. Zhang J, Goliwas KF, Wang W, Taufalele PV, Bordeleau F, Reinhart-King CA. Energetic regulation of coordinated leader-follower dynamics during collective invasion of breast cancer cells. Proc Natl Acad Sci U S A 2019, 116(16): 7867–7872.

8. Riahi R, Sun J, Wang S, Long M, Zhang DD, Wong PK. Notch1-Dll4 signalling and mechanical force regulate leader cell formation during collective cell migration. Nat Commun 2015, 6: 6556.

9. Huarte M. The emerging role of lncRNAs in cancer. Nat Med 2015, 21(11): 1253–1261.

10. Esteller M. Non-coding RNAs in human disease. Nat Rev Genet 2011, 12(12): 861–874.

11. Statello L, Guo C-J, Chen L-L, Huarte M. Gene regulation by long non-coding RNAs and its biological functions. Nature Reviews Molecular Cell Biology 2021, 22(2): 96–118.

12. Iyer MK, Niknafs YS, Malik R, Singhal U, Sahu A, Hosono Y, et al. The landscape of long noncoding RNAs in the human transcriptome. Nature Genetics 2015, 47(3): 199–208.

13. Guttman M, Rinn JL. Modular regulatory principles of large non-coding RNAs. Nature 2012, 482(7385): 339–346.

14. Slack FJ, Chinnaiyan AM. The Role of Non-coding RNAs in Oncology. Cell 2019, 179(5): 1033–1055.

15. Schmitt AM, Chang HY. Long Noncoding RNAs in Cancer Pathways. Cancer Cell 2016, 29(4): 452–463.

16. Arun G, Aggarwal D, Spector DL. MALAT1 Long Non-Coding RNA: Functional Implications. Noncoding RNA 2020, 6(2).

17. Zhan Y, Du L, Wang L, Jiang X, Zhang S, Li J, et al. Expression signatures of exosomal long non-coding RNAs in urine serve as novel non-invasive biomarkers for diagnosis and recurrence prediction of bladder cancer. Mol Cancer 2018, 17(1): 142.

18. Arun G, Diermeier S, Akerman M, Chang KC, Wilkinson JE, Hearn S, et al. Differentiation of mammary tumors and reduction in metastasis upon Malat1 lncRNA loss. Genes Dev 2016, 30(1): 34–51.

19. Gutschner T, Hammerle M, Eissmann M, Hsu J, Kim Y, Hung G, et al. The noncoding RNA MALAT1 is a critical regulator of the metastasis phenotype of lung cancer cells. Cancer Res 2013, 73(3): 1180–1189.

20. Sun R, Qin C, Jiang B, Fang S, Pan X, Peng L, et al. Down-regulation of MALAT1 inhibits cervical cancer cell invasion and metastasis by inhibition of epithelial-mesenchymal transition. Mol Biosyst 2016, 12(3): 952–962.

21. Kwok ZH, Roche V, Chew XH, Fadieieva A, Tay Y. A non-canonical tumor suppressive role for the long non-coding RNA MALAT1 in colon and breast cancers. Int J Cancer 2018, 143(3): 668–678.

22. Han Y, Wu Z, Wu T, Huang Y, Cheng Z, Li X, et al. Tumor-suppressive function of long noncoding RNA MALAT1 in glioma cells by downregulation of MMP2 and inactivation of ERK/MAPK signaling. Cell Death Dis 2016, 7: e2123.

23. Kim J, Piao HL, Kim BJ, Yao F, Han Z, Wang Y, et al. Long noncoding RNA MALAT1 suppresses breast cancer metastasis. Nat Genet 2018, 50(12): 1705–1715.

24. Xu S, Sui S, Zhang J, Bai N, Shi Q, Zhang G, et al. Downregulation of long noncoding RNA MALAT1 induces epithelial-to-mesenchymal transition via the PI3K-AKT pathway in breast cancer. Int J Clin Exp Pathol 2015, 8(5): 4881–4891.

25. Tutucci E, Vera M, Biswas J, Garcia J, Parker R, Singer RH. An improved MS2 system for accurate reporting of the mRNA life cycle. Nat Methods 2017.

26. Cawte AD, Unrau PJ, Rueda DS. Live cell imaging of single RNA molecules with fluorogenic Mango II arrays. Nat Commun 2020, 11(1): 1283.

27. Kozyrska K, Pilia G, Vishwakarma M, Wagstaff L, Goschorska M, Cirillo S, et al. p53 directs leader cell behavior, migration, and clearance during epithelial repair. Science 2022, 375(6581): eabl8876.

28. Chrisafis G, Wang T, Moissoglu K, Gasparski AN, Ng Y, Weigert R, et al. Collective cancer cell invasion requires RNA accumulation at the invasive front. Proc Natl Acad Sci U S A 2020, 117(44): 27423–27434.

29. Young AP, Jackson DJ, Wyeth RC. A technical review and guide to RNA fluorescence in situ hybridization. PeerJ 2020, 8: e8806.

30. Ozsolak F, Milos PM. RNA sequencing: advances, challenges and opportunities. Nat Rev Genet 2011, 12(2): 87–98.

31. Santangelo PJ, Nix B, Tsourkas A, Bao G. Dual FRET molecular beacons for mRNA detection in living cells. Nucleic Acids Res 2004, 32(6): e57.

32. Wan Y, Zhu N, Lu Y, Wong PK. DNA Transformer for Visualizing Endogenous RNA Dynamics in Live Cells. Anal Chem 2019, 91(4): 2626–2633.

33. Riahi R, Dean Z, Wu TH, Teitell MA, Chiou PY, Zhang DD, et al. Detection of mRNA in living cells by double-stranded locked nucleic acid probes. Analyst 2013, 138: 4777–4785.

34. Xu J, Lamouille S, Derynck R. TGF-beta-induced epithelial to mesenchymal transition. Cell Res 2009, 19(2): 156–172.

35. Riahi R, Yang YL, Zhang DD, Wong PK. Advances in wound-healing assays for probing collective cell migration. J Lab Autom 2012, 17(1): 59–65.

36. Bocci F, Tripathi SC, Vilchez Mercedes SA, George JT, Casabar JP, Wong PK, et al. NRF2 activates a partial epithelial-mesenchymal transition and is maximally present in a hybrid epithelial/mesenchymal phenotype. Integr Biol (Camb) 2019, 11(6): 251–263.

37. Vilchez Mercedes SA, Bocci F, Zhu N, Levine H, Onuchic JN, Jolly MK, et al. Nrf2 modulates the hybrid epithelial/mesenchymal phenotype and Notch signaling during collective cancer migration. Frontiers in Molecular Biosciences 2022, 9: 807324.

38. Park HY, Lim H, Yoon YJ, Follenzi A, Nwokafor C, Lopez-Jones M, et al. Visualization of dynamics of single endogenous mRNA labeled in live mouse. Science 2014, 343(6169): 422–424.

39. Vilchez Mercedes SA, Eder I, Ahmed M, Zhu N, Wong PK. Optimizing locked nucleic acid modification in double-stranded biosensors for live single cell analysis. Analyst 2022, 147(4): 722–733.

40. LeSavage BL, Suhar RA, Broguiere N, Lutolf MP, Heilshorn SC. Nextgeneration cancer organoids. Nat Mater 2022, 21(2): 143–159.

41. Yang YL, Jamilpour N, Yao BY, Dean ZS, Riahi R, Wong PK. Probing Leader Cells in Endothelial Collective Migration by Plasma Lithography Geometric Confinement. Sci Rep-Uk 2016, 6: 22707.

42. Chen M, Xia Z, Chen C, Hu W, Yuan Y. LncRNA MALAT1 promotes epithelial-to-mesenchymal transition of esophageal cancer through Ezh2-Notch1 signaling pathway. Anticancer Drugs 2018, 29(8): 767–773.

43. Zhao C, Ling X, Xia Y, Yan B, Guan Q. The m6A methyltransferase METTL3 controls epithelial-mesenchymal transition, migration and invasion of breast cancer through the MALAT1/miR-26b/HMGA2 axis. Cancer Cell Int 2021, 21(1): 441.

44. Sun Z, Ou C, Liu J, Chen C, Zhou Q, Yang S, et al. YAP1-induced MALAT1 promotes epithelial-mesenchymal transition and angiogenesis by sponging miR-126-5p in colorectal cancer. Oncogene 2019, 38(14): 2627–2644.

45. Zhou Y, Shan T, Ding W, Hua Z, Shen Y, Lu Z, et al. Study on mechanism about long noncoding RNA MALAT1 affecting pancreatic cancer by regulating Hippo-YAP signaling. J Cell Physiol 2018, 233(8): 5805–5814.

46. Zuker M. Mfold web server for nucleic acid folding and hybridization prediction. Nucleic Acids Research 2003, 31(13): 3406–3415.

47. Jaqaman K, Loerke D, Mettlen M, Kuwata H, Grinstein S, Schmid SL, et al. Robust single-particle tracking in live-cell time-lapse sequences. Nature Methods 2008, 5(8): 695–702.

48. Tinevez J-Y, Perry N, Schindelin J, Hoopes GM, Reynolds GD, Laplantine E, et al. TrackMate: An open and extensible platform for single-particle tracking. Methods 2017, 115: 80–90.

