## Supplementary Information for "Long Noncoding RNA MALAT1 is Dynamically Regulated in Leader Cells during Collective Cancer Invasion"

Ninghao Zhu,^1^ Mona Ahmed,^1^ Joseph C. Liao,^2^ and Pak Kin Wong^1,3^

^1^Department of Biomedical Engineering, The Pennsylvania State University, University Park, PA, USA

^2^Department of Urology, Stanford University School of Medicine, Stanford, California, USA.

^3^Department of Mechanical Engineering and Department of Surgery, The Pennsylvania State University, University Park, PA, USA

Table of Contents:

Supplementary Tables 1 to 3

Supplementary Figures 1 to 11

**Supplementary Table 1.** The sequence of β-actin double stranded LNA probes and synthetic DNA targets (+N represents LNA monomer).

| **Sequence Name** | **5' Modification** | **Sequence (5’ to 3’)** | **3' Modification** |
| --- | --- | --- | --- |
| β-actin Donor Probe | ATTO488 | +AG+CC+AG+GT+CC+AG+AC+GC+AG+GA | None |
| β-actin Donor Quencher | None | GGACCTGGCT | Iowa Black FQ |
| β-actin Donor Target | None | TCCTGCGTCTGGACCTGGCT | None |
| β-actin Acceptor Probe | None | +GA+GG+TA+GT+CA+GT+CA+GG+TC+CC | Cy3 |
| β-actin Acceptor Quencher | Iowa Black FQ | GGGACCTGAC | None |
| β-actin Acceptor Target | None | GGGACCTGACTGACTACCTC | None |
| β-actin Full Target | None | TCCTGCGTCTGGACCTGGCTGGCCGGGACCTGACTGACTACCTC | None |

**Supplementary Table 2.** The sequence of MALAT1 double stranded LNA probes and synthetic DNA targets (+N represents LNA monomer).

| **Sequence Name** | **5' Modification** | **Sequence (5’ to 3’)** | **3' Modification** |
| --- | --- | --- | --- |
| MALAT1Donor Probe | ATTO488 | +AT+GT+TC+CC+AC+CC+AG+CA+TT +AC | None |
| MALAT1Donor Quencher | None | GTGGGAACAT | Iowa Black FQ |
| MALAT1Donor Target | None | GTAATGCTGGGTGGGAACAT | None |
| MALAT1Acceptor Probe | None | +AT+CT+TC+TC+CA+GT+CT+AC+AA+GT | Cy3 |
| MALAT1Acceptor Quencher | Iowa Black FQ | ACTTGTAGAC | None |
| MALAT1Acceptor Target | None | ACTTGTAGACTGGAGAAGAT | None |
| MALAT1Full Target | None | GTAATGCTGGGTGGGAACATGTAACTTGTAGACTGGAGAAGAT | None |

**Supplementary Table 3.** The sequence of UCA1 double stranded LNA probes and synthetic DNA targets (+N represents LNA monomers).

| **Sequence Name** | **5' Modification** | **Sequence (5’ to 3’)** | **3' Modification** |
| --- | --- | --- | --- |
| UCA1 Donor Probe | FAM | +TC+TG+GA+AT+GG+TG+AA+CC+CA+AT | None |
| UCA1 Donor Quencher | None | CCATTCCAGA | Iowa Black FQ |
| UCA1 Donor Target | None | ATTGGGTTCACCATTCCAGA | None |
| UCA1 Acceptor Probe | None | +GC+TG+TC+TG+AT+GG+GC+AT+GG+CT | TexRd-XN |
| UCA1 Acceptor Quencher | Iowa Black RQ | AGCCATGCCC | None |
| UCA1 Acceptor Target | None | AGCCATGCCCATCAGACAGC | None |
| UCA1 Full Target | None | ATTGGGTTCACCATTCCAGAATAAAGCCATGCCCATCAGACAGC | None |

**Supplementary Figures**

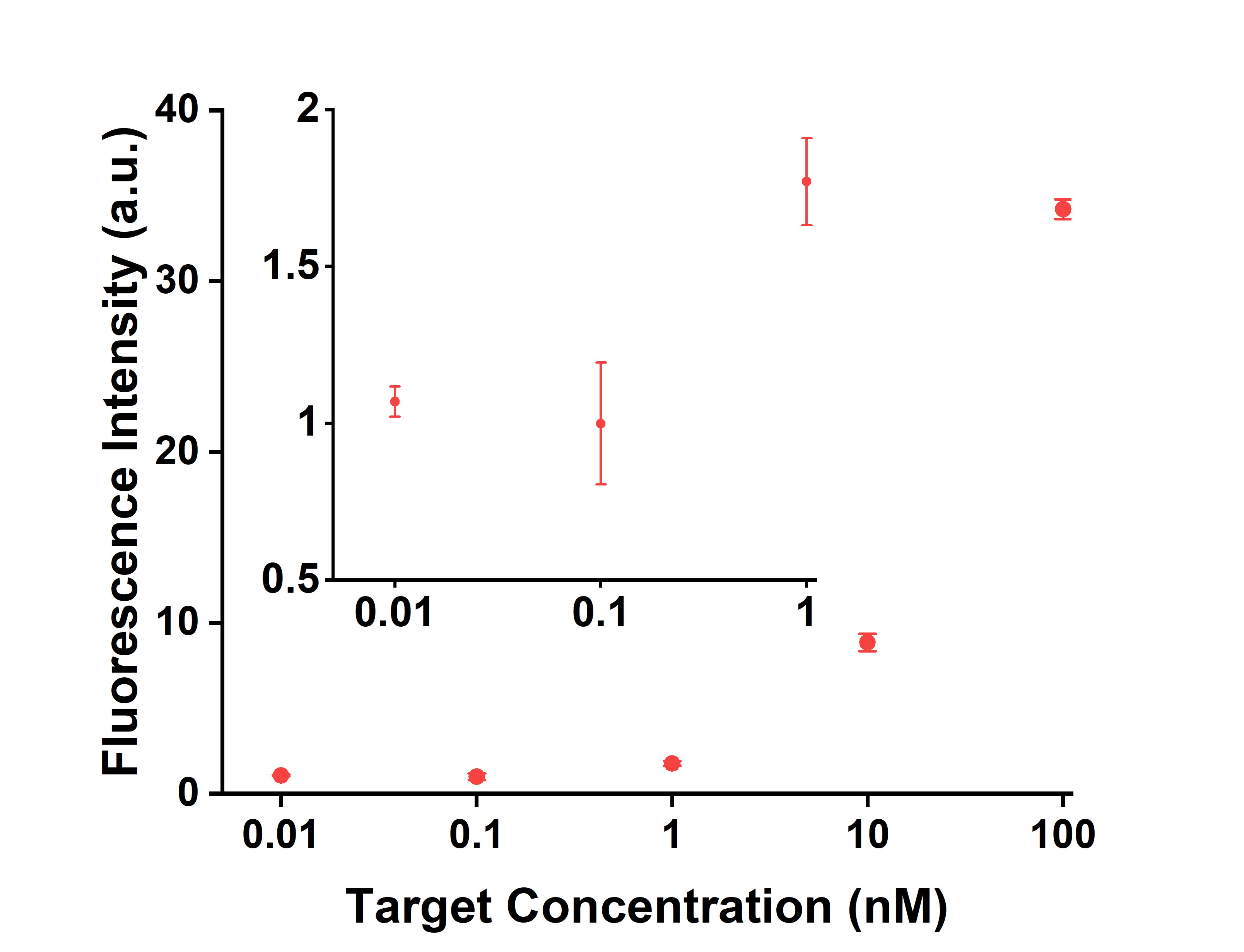

**Fig. S1 |** Characterization of the dual dsLNA probe for detecting a synthetic target sequence (n = 3).

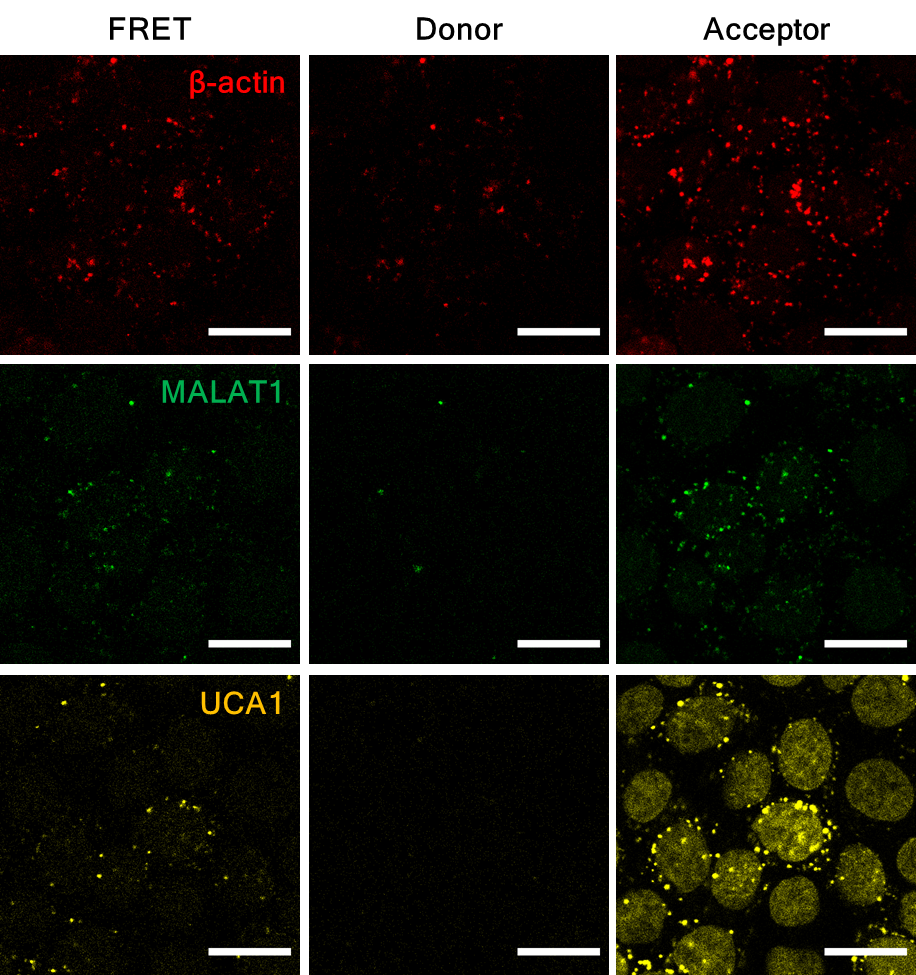

**Fig. S2 |** Confocal images of β-actin, MALAT1, and UCA1 probes in FRET, donor, and acceptor channels for RNA detection in live bladder cancer cells (5637). Scale bars, 50 μm. Images are representative of eight experiments.

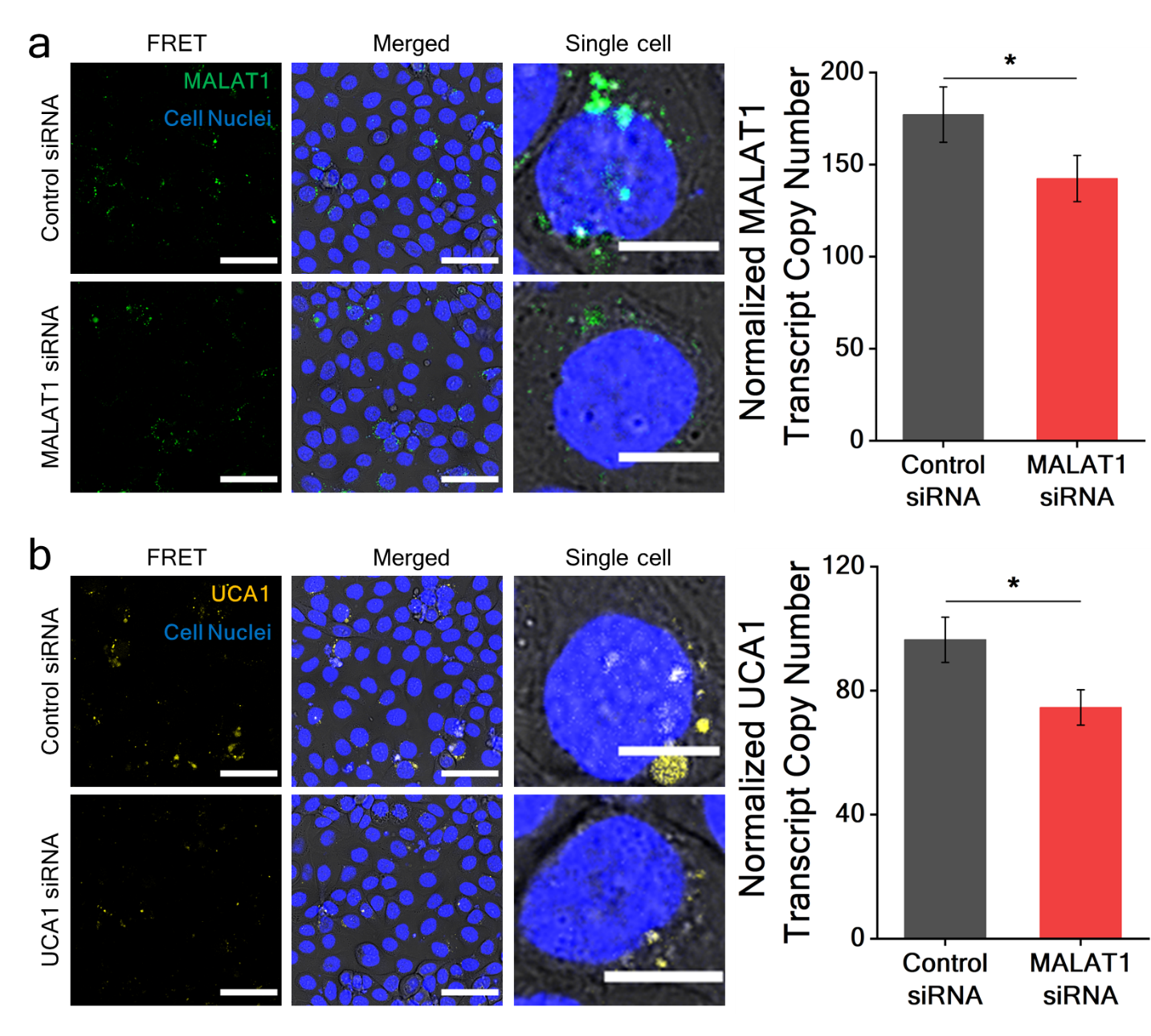

**Fig. S3 |** Quantification of lncRNA transcripts in live cells. **a**, Detection of MALAT1 by confocal microscopy in live cells with MALAT1 siRNA and control siRNA. **b**, Detection of UCA1 by confocal microscopy in live cells with UCA1 siRNA and control siRNA. Scale bars, 50 μm (FRET, merged channels) and 10 μm (single cell channel). Images are representative of four experiments. Student’s t-test was used to compare transcript numbers between control siRNA and lncRNA siRNA treatment (n ≥ 9, * p < 0.05).

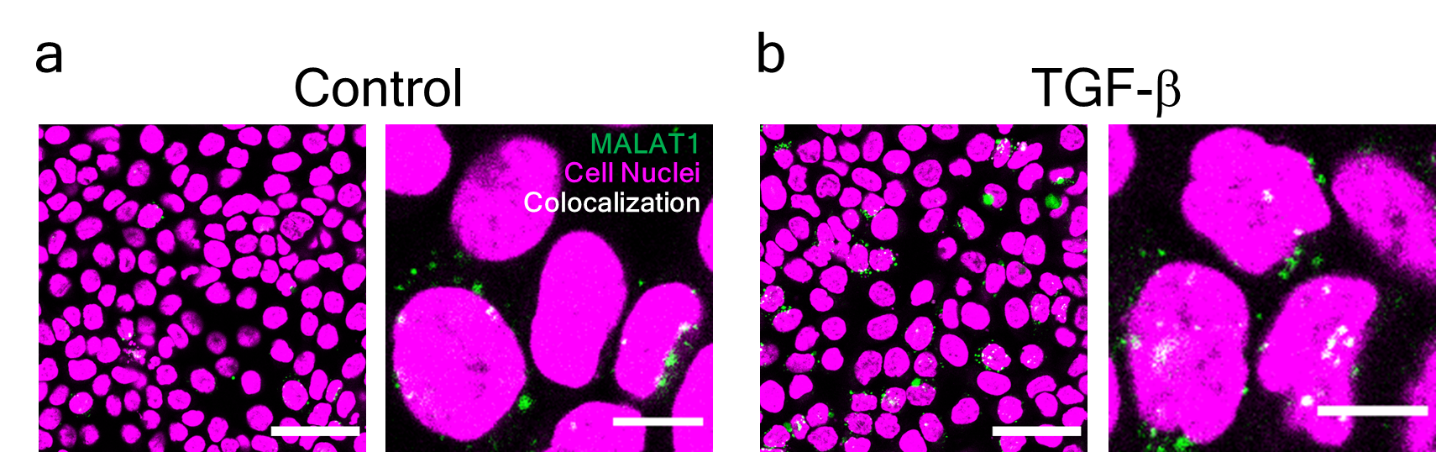

**Fig. S4 |** Confocal images of MALAT1 dual probe and cell nucleus counterstain in live cells (5637) treated with (**a**) buffer control and (**b**) TGF-β. Images are representative of five experiments.

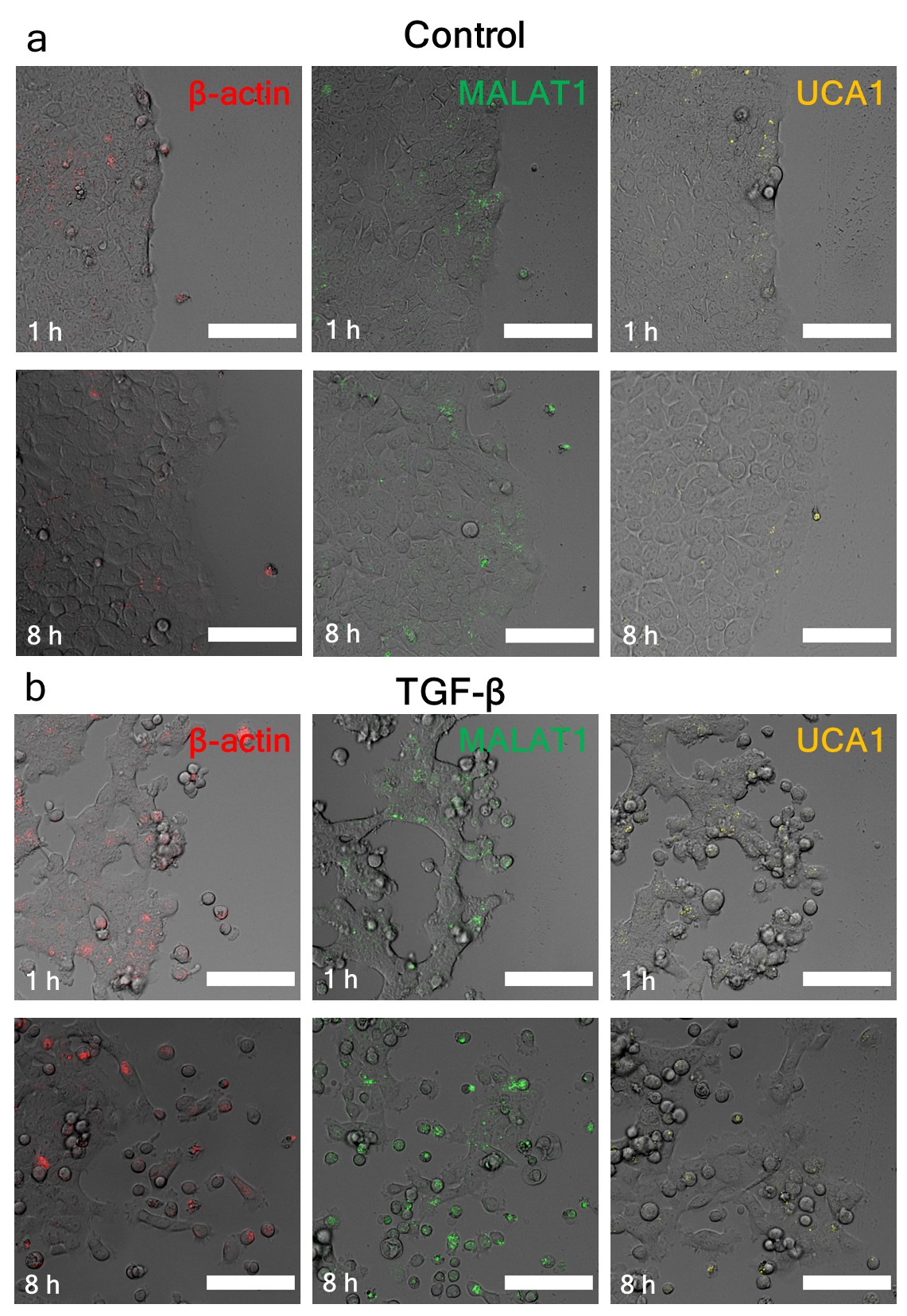

**Fig. S5 |** MALAT1 expression in the scratch cell migration assay. **a**, Confocal images of cells near the migrating front at 1 hour and 8 hours. **b**, Confocal images of cells near the migrating front at 1 hour and 8 hours. The cells were treated with TGF-β. Scale bars, 100 µm. Images are representative of three experiments.

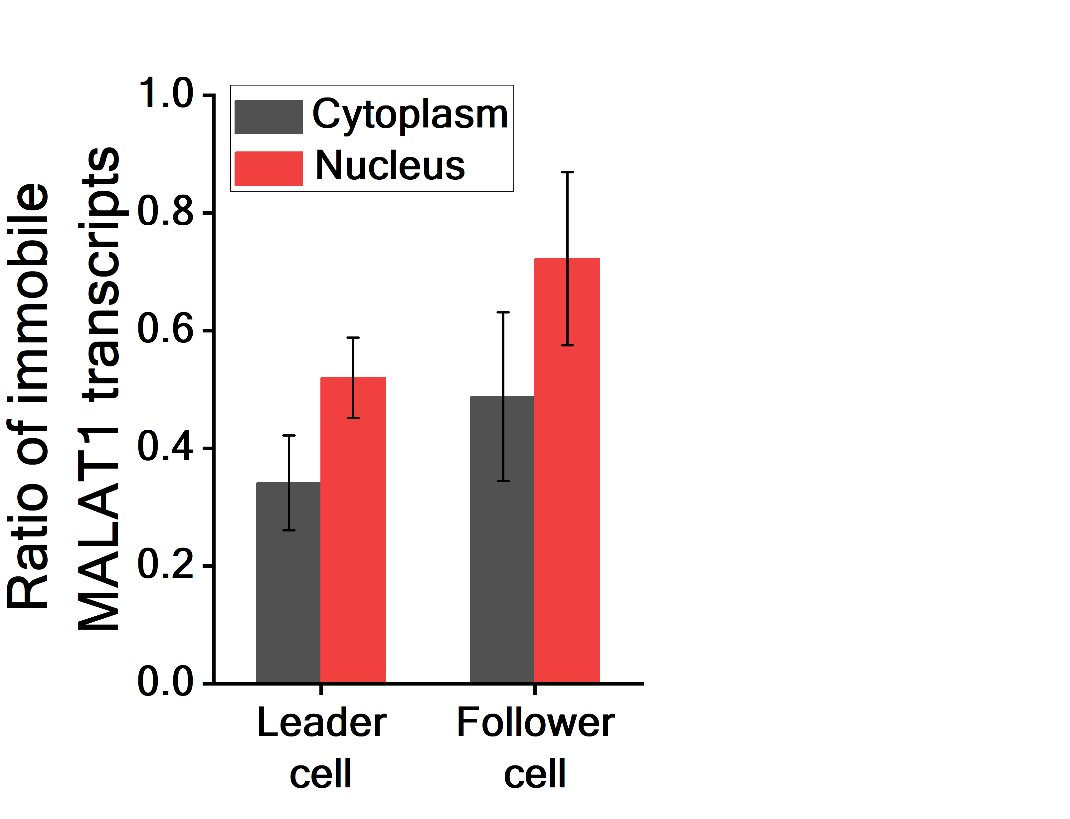

**Fig. S6 |** The portion of immobile MALAT1 transcripts (diffusivity < 0.005 μm^2^/s) in leader cells and follower cells. Two-way ANOVA did not show a significance between the values (n = 5 for leader cells and n = 3 for follower cells).

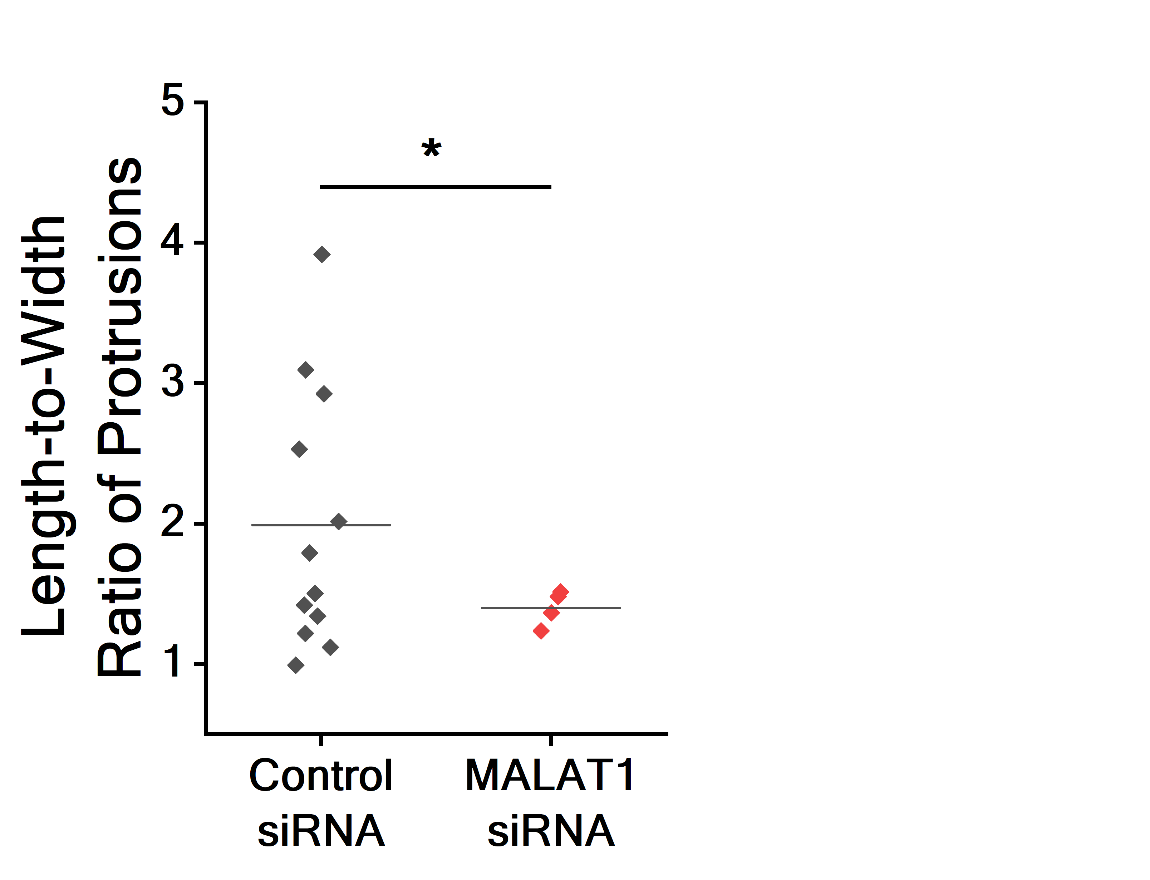

**Fig. S7 |** The length-to-width ratio of the protrusions from cancer spheroids treated with MALAT1 siRNA and control siRNA on day 3. Student’s t-test was used to compare the length-to-width ratio in the MALAT1 knockdown spheroids and control spheroids. *, p < 0.05 (n = 4 MALAT1 siRNA and n = 12 for control siRNA).

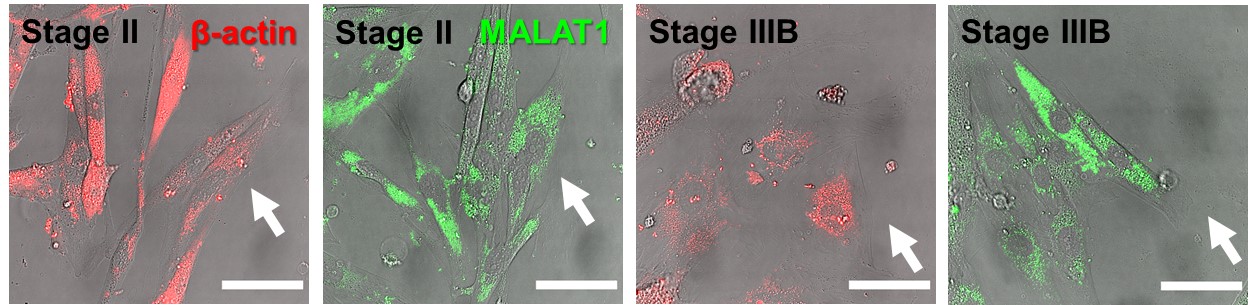

**Fig. S8 |** β-actin mRNA and MALAT1 expression (white arrows) of human dissociated bladder cancer cells in the scratch cell migration assay. Scale bars, 50 μm. Images are representative of 10 leader cells.

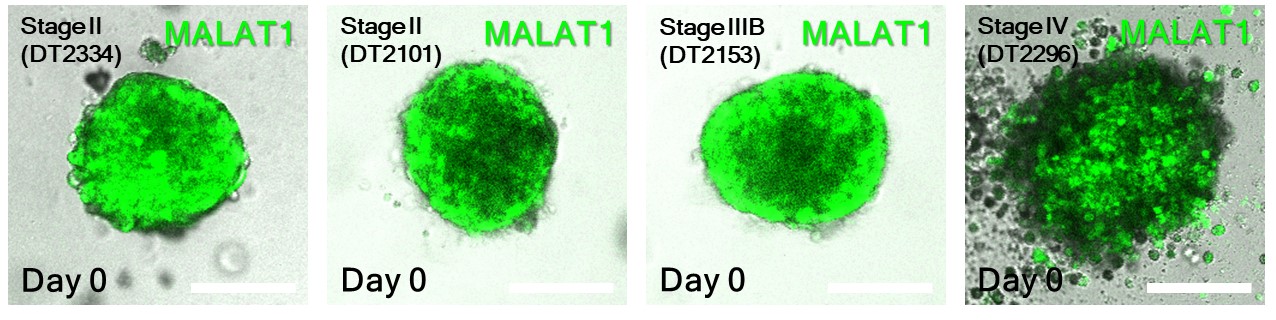

**Fig. S9 |** MALAT1 expression in human bladder tumor organoids on day 0. Scale bars, 200 µm.

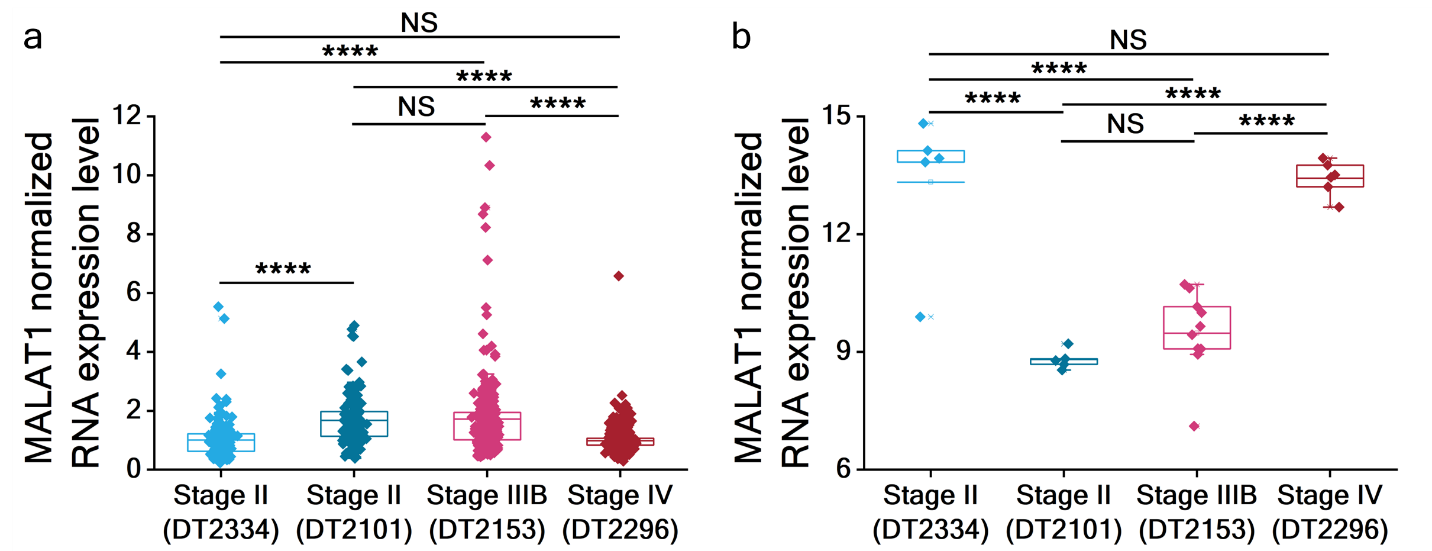

**Fig. S10 | a,** Normalized MALAT1 expression in detached cells. One-way ANOVA followed by Tukey’s post hoc test was used to compare the MALAT1 expression level in detached cells of different human samples, NS, not significant, ****, p < 0.0001 (n = 178, 128, 270, and 642 for DT2334, DT2101, DT2153, and DT2296). **b**, Normalized lncRNA MALAT1 expression in the bulk spheroids. One-way ANOVA followed by Tukey’s post hoc test was used to compare the MALAT1 expression level in leader cells of different human samples, NS, not significant, ****, p < 0.0001 (n = 5, 5, 10, 6 for DT2334, DT2101, DT2153 and DT2296).

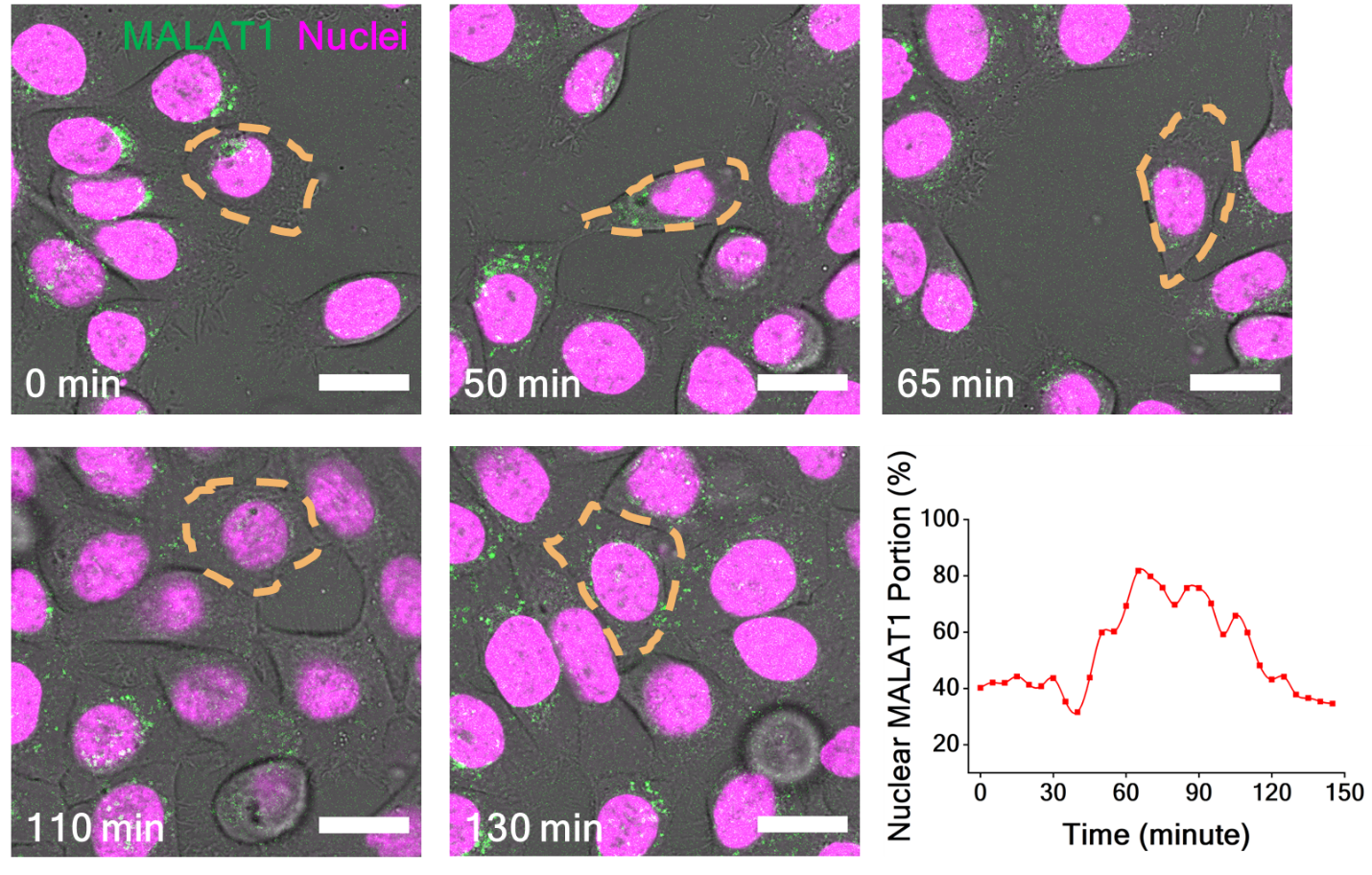

**Fig. S11 |** Dynamics of nuclear MALAT1 in leader cells. The leader cell is outlined with yellow dashes. Images are representative of five leader cells. Scale bars, 20 µm.

**Supplementary Video Caption**

**Supplementary Video 1.** 3D rendering and segmentation of an invading cancer spheroid in the 3D invasion assay. MALAT1 expression is indicated by the red signal. The invading sprouts and detached cells are highlighted by the green contours.

**Supplementary Video 2.** Single molecule tracking of MALAT1 transcripts in a leader cell in 301 seconds. MALAT1 transcripts are indicated by the green dots, and the cell nucleus is marked with the blue signal. Scale bar, 10 μm.

**Supplementary Video 3.** Single molecule tracking of MALAT1 transcripts in a follower cell in 301 seconds. MALAT1 transcripts are indicated by the green dots, and the cell nucleus is marked with the blue signal. Scale bar, 10 μm.

**Supplementary Video 4.** MALAT1 transcript tracking during the closure of the scratch cell migration assay. MALAT1 transcripts are indicated by the green dots, and the cell nucleus is marked with the blue signal. Scale bars, 50 μm.

**Supplementary Video 5.** MALAT1 transcript tracking during the closure of the scratch cell migration assay. MALAT1 transcripts are indicated by the green dots, and the cell nucleus is marked with the blue signal. The leader cell is indicated by the white asterisk. Scale bars, 50 μm.
